# Development of a Confinable Gene-Drive System in the Human Disease Vector, *Aedes aegypti*

**DOI:** 10.1101/645440

**Authors:** Ming Li, Ting Yang, Nikolay P. Kandul, Michelle Bui, Stephanie Gamez, Robyn Raban, Jared Bennett, Héctor M. Sánchez C., Gregory C. Lanzaro, Hanno Schmidt, Yoosook Lee, John M. Marshall, Omar S. Akbari

## Abstract

*Aedes aegypti*, the principal mosquito vector for many arboviruses that causes yellow fever, dengue, Zika, and chikungunya, increasingly infects millions of people every year. With an escalating burden of infections and the relative failure of traditional control methods, the development of innovative control measures has become of paramount importance. The use of gene drives has recently sparked significant enthusiasm for the genetic control of mosquito populations, however no such system has been developed in *Ae. aegypti*. To fill this void and demonstrate efficacy in *Ae. aegypti,* here we develop several CRISPR-based split-gene drives for use in this vector. With cleavage rates up to 100% and transmission rates as high as 94%, mathematical models predict that these systems could spread anti-pathogen effector genes into wild *Ae. aegypti* populations in a safe, confinable and reversible manner appropriate for field trials and effective for controlling disease. These findings could expedite the development of effector-linked gene drives that could safely control wild populations of *Ae. aegypti* to combat local pathogen transmission.

**Significance Statement:** *Ae. aegypti* is a globally distributed arbovirus vector spreading deadly pathogens to millions of people annually. Current control methods are inadequate and therefore new technologies need to be innovated and implemented. With the aim of providing new tools for controlling this pest, here we engineered and tested several split gene drives in this species. These drives functioned at very high efficiency and may provide a tool to fill the void in controlling this vector. Taken together, our results provide compelling path forward for the feasibility of future effector-linked split-drive technologies that can contribute to the safe, sustained control and potentially the elimination of pathogens transmitted by this species.

## Introduction

Due to the high annual incidence of vector-borne disease, the associated economic burdens, and the lack of effective vaccines, interest in the innovation of novel population-control methods to prevent pathogen transmission is increasing. *Aedes aegypti* is the principal mosquito vector of several arboviruses, including yellow fever, chikungunya, dengue, and Zika (1). This mosquito alone places roughly half of the world’s population at risk of acquiring vector-borne diseases while causing an estimated 390 million dengue infections annually (2). Traditional mosquito control strategies, including long-lasting insecticide-treated bed nets (LLINs), chemical insecticides, and environmental management (3), have significantly reduced mosquito-borne disease burdens; however these interventions are largely unsustainable due to the development of both genetic and behavioral vector resistance that decrease their efficacy (4), high costs, and the need for repeated applications with limited spatial impact. Furthermore, chemical interventions, a fundamental tool for vector control, negatively affect non-target beneficial organisms, such as pollinators. Therefore, there is a pressing global need for alternative safe and effective approaches to control mosquito disease vectors.

Species-specific mosquito population suppression technologies, such as the *Wolbachia* incompatible insect technique (IIT) and the release of dominant lethal gene (RIDL), are currently being implemented on a small-scale in the field to control populations of *Ae. aegypti* (*5–7*). While these technologies significantly improve *Ae. aegypti* control capabilities, they require continuous inundation, which is laborious and cost prohibitive to many areas with the largest vector-borne disease burdens. In addition to IIT, *Wolbachia* has also been used for population replacement in various locations (8–10), as certain strains can spread to fixation while greatly reducing the vector’s susceptibility to dengue virus (11). However, the literature suggests the effectiveness of different *Wolbachia* strains varies so far as to even enhance vector competence in some contexts, which may lead to a reevaluation of this technique (12–18). Therefore, there has been renewed interest in developing novel species-specific and cost-effective genetic technologies that provide long-lasting disease reduction with limited effort.

One innovative technology, first articulated by Austin Burt in 2003 (19), utilizes homing-based gene-drive technologies to expedite the elimination and eradication of vector-borne diseases (20–25). Conceptually, these drives function by exploiting the organism’s innate DNA repair machinery to copy or “home” themselves into a target genomic location prior to meiosis in the germline. This process converts wild-type alleles into drive alleles in heterozygotes, thereby forcing super-mendelian inheritance of the drive into subsequent generations, irrespective of the fitness cost to the organism. Theoretically, this inheritance scheme can spread the drive and any linked anti-pathogen “cargo genes,” to fixation in a population in a short time frame (26–28), even with modest introduction frequencies (19). Importantly, if these drives are linked with effective anti-pathogen cargo genes, they could deliver efficient and cost-effective vector control (19). The drawback to this scheme came from the difficulty in engineering homing drives with various endonucleases, however the recent CRISPR revolution has streamlined their development (20, 21). In fact, proof-of-concept CRISPR-based homing gene drives have recently been developed in several organisms including yeast (29–32), flies (33–38), mice (39), and two malaria vector species, *Anopheles gambiae* (*40–42*) and *Anopheles stephensi* (43).

Unfortunately, a gene-drive system has yet to be developed in *Ae. aegypti*, the lack of which is principally due to the absence of tools necessary to engineer drives. In response to these deficiencies, here we molecularly characterized drive components and then evaluated and optimized these components in several drive systems in *Ae aegypti*. As a safety precaution, we engineered “split drives” with a molecularly unlinked endonuclease, thereby allowing the user to spatially and temporally confine the drive (20, 21, 44, 45). To improve drive efficiency, we targeted a highly conserved region in a phenotypic gene with cleavage rates, indicating biallelic cutting, of 100% and homing rates, indicating the super-Mendelian transmission rate of the drive, as high as 94%. In addition to demonstrating drive efficiency, we performed mathematical modeling that suggested split-drives are particularly valuable in gene-drive development since: *i)* several consecutive releases of male mosquitoes harboring the split-drive and anti-pathogen cargo, at an achievable 1:1 ratio with the wild population, spread disease refractoriness into wild populations, *ii)* the split-drive does not significantly spread into neighboring populations, and *iii)* dissociation of the unlinked endonuclease and guide RNA components, in addition to the fitness costs associated with the split drive and cargo, cause the drive to be eliminated from the population on a timescale that could accommodate local arbovirus elimination. These highly desirable features could enable safe testing of a split-drive system in the field prior to the release of a more invasive, self-propagating, linked-drive system and could lead to the development of new technologies to prevent vector-borne disease transmission.

## Results

### Rational target site selection for the drive

To develop a binary Cas9 and guide RNA (gRNA) approach that could be leveraged for the development of gene drives, we built on our previous work that generated *Ae. aegypti* strains expressing the Cas9 endonuclease in both germline and somatic cells and demonstrated their robust cleavage and improved homology directed repair (HDR) efficiencies (46). We targeted an easily screenable gene with consistent and viable disrupted phenotype, the *white* gene (AAEL016999), which is an ATP-binding cassette (ABC) transporter (46–48). To locate a highly conserved target sequence in *white*, we bioinformatically compared the whole-genome sequences of 133 *Ae. aegypti* mosquitoes sampled throughout California, Florida, Mexico, and South Africa (Fig. S1, Table S1) (49). From this analysis, we discovered a putative gRNA target sequence in exon 3 of *white* (*gRNA^w^*) that was 100% conserved in these wild populations (Fig. S2A–B). To target this highly conserved sequence and determine cleavage efficiency at this site, we synthesized and injected gRNAs into embryos derived from *exu-Cas9* (*Exuperantia*, AAEL010097) females that maternally deposit Cas9. We achieved a 92.7 ± 5.8% mutation efficiency in the injected G_0_s, which is consistent with our previous work in this line (46), demonstrating that this target site is accessible to Cas9 cleavage and that this gRNA is highly active.

### Characterization of functional polymerase-III promoters

Next, we needed a promoter to ensure proper strength and timing of gRNA expression for cleavage and homing to occur in the germline. U6 non-coding small nuclear RNA (snRNA) polymerase III promoters are ideal for gRNA expression due to their nucleus-associated transcription and unmodified 5’ and 3’ regions (i.e., no 5’ cap or 3’ polyA tail); consequently, they have been used to drive gRNA expression in various species (40, 43, 50). Unfortunately, U6 promoters have not been functionally characterized in *Ae. aegypti*. There are 12 known U6 promoters in *Ae. aegypti*, therefore to characterize their usefulness for gRNA expression, we engineered 12 constructs each with a predicted U6 promoter (termed U6a-U6l from hereon) derived from *Ae. aegypti* to drive expression of *gRNA^w^* (Fig. S2C). To transiently assess promoter activity, we microinjected each construct separately into *exu-Cas9* derived embryos (46) (Fig. 1A). Of the 12 injected constructs, only four (U6a-d) induced somatic loss-of-function variegated biallelic white-eye loss-of-function phenotypes in G_0_ embryos with moderate mutation frequencies ranging from 32.5–71.3% (Fig. 1B), indicating that these four promoters can transiently express *gRNA^w^*. We then evaluated germline mutation rates by outcrossing mosaic G_0_ mosquitoes to recessive *white* mutants (*w^Δ19^/w^Δ19^*) harboring a 19-bp deletion at the *white* gRNA cut site which would lead to heritable the recessive loss-of-function phenotypes in successfully edited mosquitoes. These phenotypes were then scored in the G1 progeny with rates ranging 27.6–67.5%, indicating efficient germline editing (Fig. 1A–B).

**Fig. 1.**
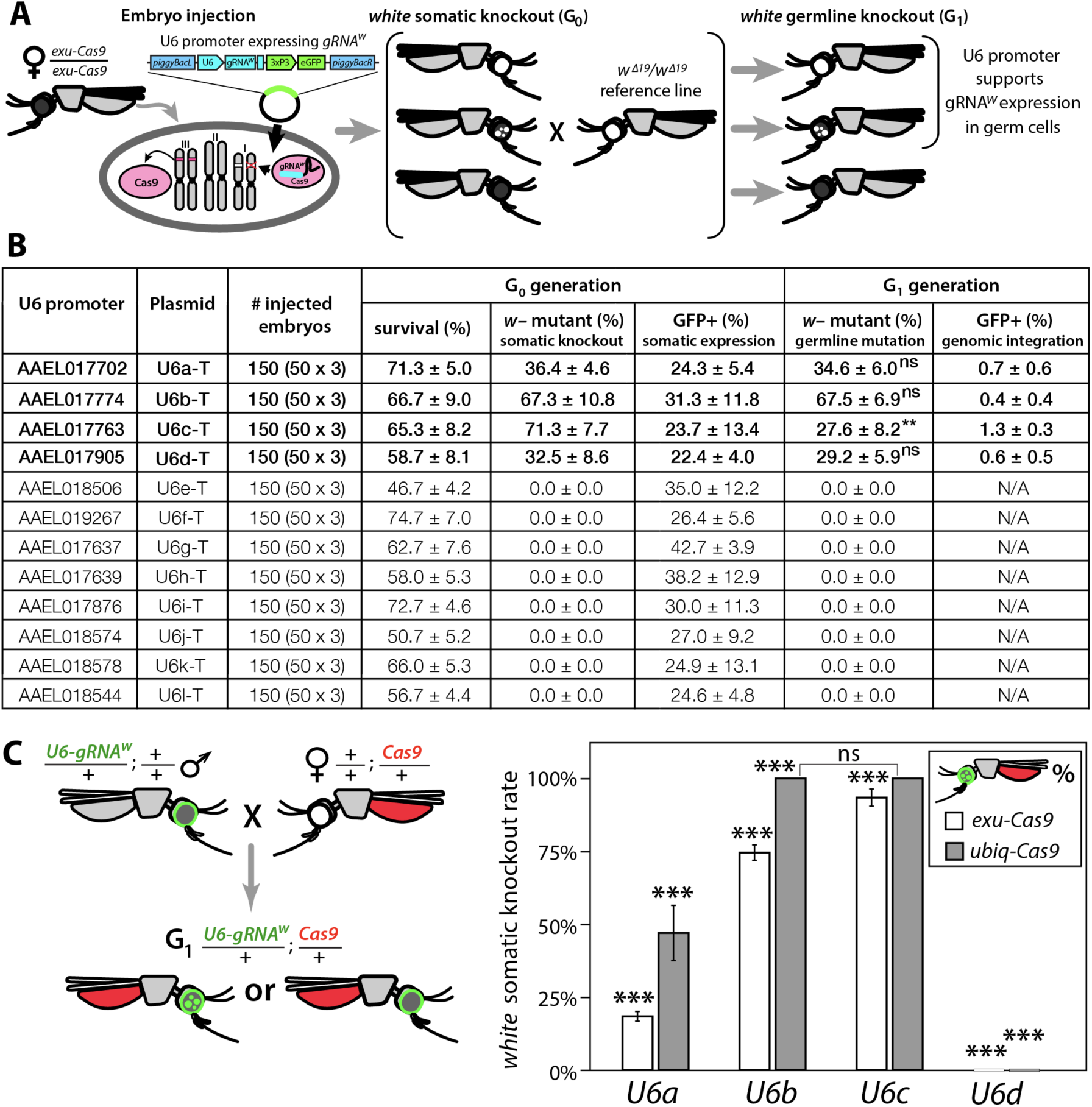
Functional identification of polymerase III promoters in *Ae. aegypti*. (A) *exu-Cas9* embryos were injected with one of 12 *piggyBac* plasmids, each utilizing a different U6 promoter expressing a guide RNA targeting the *white* eye pigmentation gene, *gRNA^w^* (Fig. S2). The frequency of somatic *white* eye phenotypes in the resulting G_0_ progeny was used to assess promoter efficiency and to confirm gRNA target site accessibility. Germline mutagenesis rates were assessed by crossing G_0_ to *white* loss-of-function (*w*^Δ*19*^/*w*^Δ*19*^) lines to determine the *white* eye phenotype frequency in G_1_ progeny. (B). Two types of *w*–knockouts were observed: complete white eyes and mosaic white eyes. Out of 12 tested U6 promoters, four U6 promoters (U6a, U6b, U6c, and U6d) induced *white* knockout phenotypes. Statistical differences between germline and somatic mutation rates were estimated by equal variance *t-*test. (C) Transgenic males harboring *piggyBac*-integrated *U6-gRNA^W^* were outcrossed to either *exu-Cas9* or *ubiq-Cas9* females (left panel), and eye phenotypes were scored in G_1_ trans-heterozygous progeny (right panel, Table S2). Statistical differences in mutation knockout rates were estimated by equal variance *t-*test. (*P* ≤ 0.05^ns^, *P* > 0.05*, *P* > 0.01**, and *P* > 0.001***)

### Development of a binary CRISPR approach

To measure cleavage and mutagenesis efficiencies of genome-integrated drive components, we developed a simple fluorescence-based system that would be compatible and orthogonal to the white mosaic readout. We developed this system by genetically encoding the four functional U6a-U6d-*gRNA^w^* transgenes by injecting these constructs into *Ae. aegypti* embryos (designated as wild type [*wt*]) and establishing one line for each construct. Transgenesis was confirmed by the enhanced green fluorescent protein (eGFP) transgenesis marker (Fig. S2C), and positives were outcrossed to *wt* strains to establish stocks. Somatic cleavage efficiencies were estimated by crossing males from each line (*U6a-U6d*-*gRNA^w^*) to females of Cas9 expression lines, *exu-Cas9* and *ubiq-Cas9* (ubiquitin L40, AAEL006511) which rely on different promoters to express Cas9 (46). The frequency of somatic *white* mutagenesis was scored in the trans-heterozygous progeny (Fig. 1C–D). Interestingly, the rates of mosaicism, an intermediate phenotype with both *wt* and knockout characteristics (i.e. black and white patchy eyes see Fig. 2), varied depending on the Cas9 and the gRNA^w^ promoters. As the timing and level of both Cas9 and gRNA expression is important for inducing mosciasm (51, 52), this result is expected. Interestingly, the combination of the *ubiq-Cas9* strain and the *U6b-gRNA^w^* and *U6c-gRNA^w^* strains induced up to 100% *white* somatic knockout frequencies, with varied rates of mosaicism, indicating that these U6 promoters are highly active and timing appropriate when encoded in the genome to induce knock-out phenotypes (Fig. 1C–D; Table S2).

**Fig. 2.**
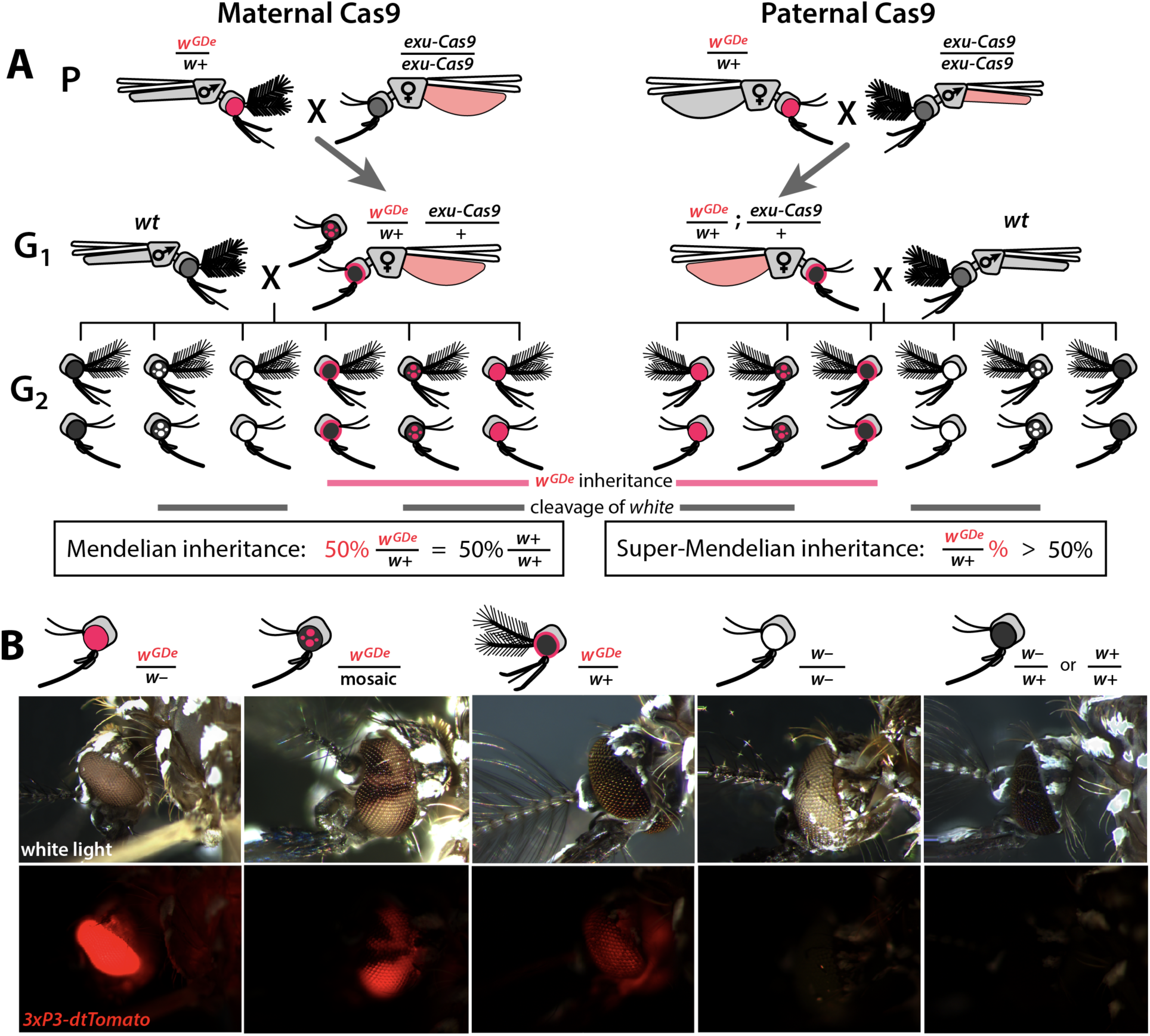
Assessing super-Mendelian inheritance of split drives. (A) Crossing scheme of the *w*^GDe^ and *exu-Cas9* parent strains (P) to generate trans-heterozygotes (G_1_), and the outcrossing of their progeny to wild-type (*wt*) mosquitoes (G_2_). *tdTomato* eye and *dsRed* abdominal expression were the *w^GDe^* and *exu-Cas9* transgene inheritance markers, respectively. *w^GDe^* transmission and *white* loss-of-function mutation rates were estimated among G_2_ progeny of trans-heterozygous female and *wt* male crosses. Super-Mendelian inheritance of *w^GDe^* occurred when the transmission rate of *w^GDe^* in G_2_ progeny was >50%, as expected by standard Mendelian inheritance. (B) Examples of G_2_ progeny eye phenotypes and corresponding genotypes.

### Engineering split drives

Given the above promising results, we next wanted to use these components to develop split gene drives. To engineer split drives, we separated the drive components into two elements: a gene-drive element (GDe) and the Cas9 endonuclease (Fig. S3). Because we did not know which U6 promoter would be optimal for homing in the germline, we engineered four GDe’s (*U6a-GDe– U6d-GDe*), each consisting of *gRNA^w^* under the control of a different functional U6 promoter (U6a–U6d) that also contained a dominant *tdTomato* marker (red fluorescent protein) for tracking and two outer homology arms immediately flanking the *gRNA^w^* target site to mediate HDR at the *white* locus (Fig. S2D, S3A). To further increase gRNA expression, the gRNA scaffold was modified slightly as previously described (53) to eliminate cryptic termination sequences. We established transgenic lines by injecting *wt* embryos with Cas9 recombinant protein pre-mixed with *gRNA*^w^ (Cas9/*gRNA*^w^) in combination with a GDe construct to serve as the template for site-specific integration via Cas9-mediated HDR (Fig. S4A). Transgenic mosquitoes, *w^GDe^/w+*, were readily identified as G_1_ larvae by their dominant *tdTomato* expression and black (*w+*) eyes, and they were then intercrossed to establish lines. Integration of the GDe’s into the *white* locus was genetically confirmed by the presence of homozygous (*w^GDe^/w^GDe^*) individuals expressing *tdTomato* with recessive white (*w–*) eyes, which was additionally confirmed by genomic sequencing of the left and right insertion boundaries (Fig. S5).

### Sex-biased inheritance of GDe

Next, we confirmed the Mendelian inheritance of each *GDe* by reciprocal outcrossing of 20 individual heterozygous (*w^GDe^/w+*) individuals to *wt* (w+/w+). The expected result was that both male and female heterozygous individuals would transmit the *w^GDe^* gene to their progeny at the Mendelian rate of 50%. Indeed, the heterozygous *w^GDe^/w+* females did meet this expectation, conferring *w^GDe^* inheritance to their progeny at normal Mendelian rates. The *w^GDe^/w+* males, however, unexpectedly displayed sex-segregated transmission of *w^GDe^* to either mostly females at 99.8% ± 0.3% (type I **♂** in Fig. S4, Table S3) or to mostly males at 99.8% ± 0.2% (type II **♂** in Fig. S4, Table S3). Although initially unexpected, *white* has been previously been descrbied as a sex-linked gene in *Ae. aegypti* (*47*), and recent genome sequencing has confirmed this linkage (54, 55) as *white* is located in close proximaty to *Nix* (AAEL022912), a dominant male-determining factor (M-factor) on chromosome I (56, 57). Therefore, depending on which chromosome the *GDe* initially integrated, the *GDe* and *Nix* were either linked, producing nearly all male progeny, which inherited both *w^GDe^* and *Nix* to exhibit male-biased inheritance, or the *GDe* was inherited separately from *Nix* (*w^GDe^/w+*) to exhibit female-biased inheritance (Fig. S4). Nevertheless, when both sexes were considered, heterozygous *w^GDe^/w+* parents transmitted *w^GDe^* to their progeny at the 50% normal Mendelian inheritance rate.

### Split drives induce super-Mendelian inheritance

To determine whether split drives could selfishly bias transmission, which could be exploited to rapidly and efficiently drive an anti-pathogen effector into a population, we performed a number of genetic crosses to determine transmission efficiencies. First, we estimated somatic cleavage efficiencies by outcrossing each *w^GDe^* strain to the *exu-Cas9* strain to generate trans-heterozygous *w^GDe^/w+*; *exu-Cas9/+* individuals. We found that rates of maternal carryover-dependent cleavage, which occurs when mothers maternally deposit high levels of Cas9 into their embryos and is indicated by the presence of mosaic phenotypes, varied depending on U6 promoter (26-100%), while the paternal carryover of Cas9 was not sufficient to induce these phenotypes (Figs. 2A, 3A; Tables S4–S5). Next, to measure homing efficiencies, we bidirectionally outcrossed G_1_ trans-heterozygotes *w^GDe^/w+*; *exu-Cas9/+* to *wt* and quantified somatic cleavage and inheritance frequencies in G_2_ progeny (Fig. 2). From these crosses, cleavage and homing efficiencies varied, most notably trans-heterozygous *w^U6c-GDe^/w+*; *exu-Cas9/+* females induced somatic mosaicism in 94.8% ± 3.9% of their progeny (Fig. 3B, Table S6). Furthermore, modest super-Mendelian inheritance was observed (71.1% ± 7.3% in G_2_ progeny) which is significantly greater than the expected 50% for normal Mendelian transmission, while no cleavage or homing was observed from trans-heterozygous G_1_ male outcrosses to *wt* (Fig. 3C; Table S7). To distinguish the contribution of maternal effects to differential drive transmission, we repeated these crosses with paternal Cas9 inheritance; however, there were no significant differences in transmission rates in the F_1_ progeny based on the parentage of the Cas9 donor (Figs. 2A, 3B–C; Tables S8–S9).

**Fig. 3.**
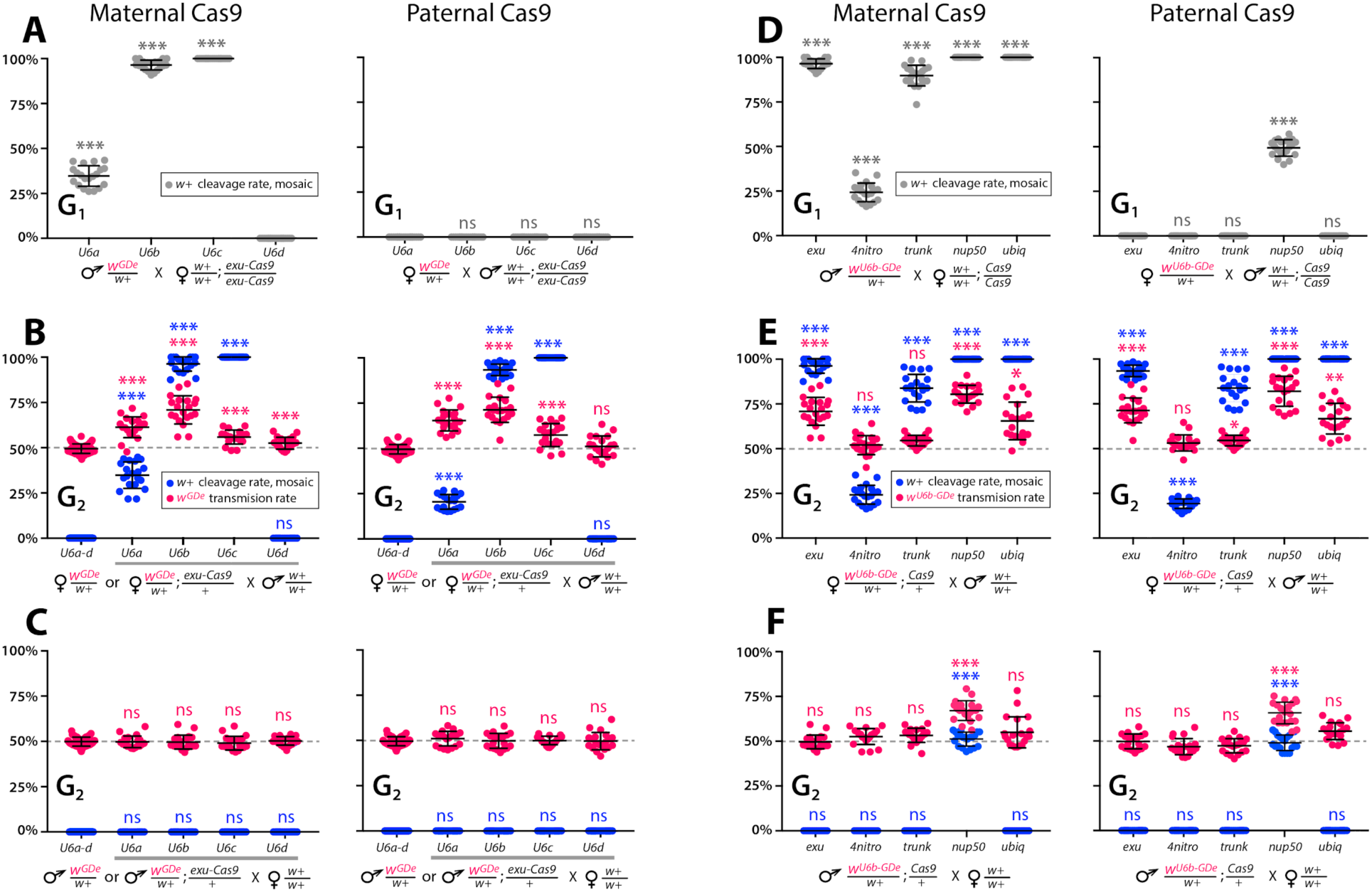
Timing and expression of drive components affect cleavage and transmission rates. We bidirectionally crossed trans-heterozygous mosquitoes that contained both components of a gene drive: the gene drive element (GDe) and the Cas9 transgene (*w^GDe^*/*w*+; *Cas9*/+). (Fig. 2A). (A–C) Four *w^GDe^/w+* lines, each with a different *gRNA^w^* U6 promoter, were crossed to the *exu-Cas9* strain to compare cleavage and homing activity. (A) Maternally deposited Cas9 protein induced *white* cleavage in G_1_ trans-heterozygotes harboring w*^U6a-GDe^, w^U6b-GDe^*, and *w^U6c-GDe^* but not *w^U6d-GDe^*. (B) In comparison to *w^GDe^/w+*, trans-heterozygous *w^GDe^/w+; exu-Cas9/+* females, (C) but not males, exhibited super-Mendelian transmission of *w^GDe^*. The *w^U6b-GDe^/w+; exu-Cas9/+* females transmitted *w^U6b-GDe^* to 71.1% ± 7.3% of G_2_ progeny. (D–F) Five lines expressing Cas9 from different promoters were crossed to *w^U6b-GDe^/w+*. (D) Maternally deposited Cas9 resulted in *white* cleavage in G_1_ trans-heterozygotes. (E) Three out of five tested Cas9 lines, *exu-Cas9, nup50-Cas9*, and *ubiq-Cas9*, induced super-Mendelian transmission of *w^GDe^* by trans-heterozygous females. The *w^U6b-GDe^/w+; nup50-Cas9/+* females transmitted *w^U6b-GDe^* to 81.3% ± 6.9% of G_2_ progeny. (F) All trans-heterozygous males transmitted *w^GDe^* following a regular Mendelian inheritance except for the *w^U6b-GDe^/w+; nup50-Cas9/+* males that induced *white* cleavage in 51.1% ± 3.9% and transmitted the *w^U6b-GDe^* allele to 66.9% ± 5.4% of G_2_ progeny. Point plots show the average ± standard deviation (SD) over 20 data points. Grey dotted line indicates standard Mendelian inheritance rates. Statistical significance was estimated using an equal variance *t-*test. (*P* ≥ 0.05^ns^, *P* < 0.05*, *P* < 0.01**, and *P* < 0.001***).

### Cas9-associated variability in homing efficiency

Given the modest homing efficiencies in females, and the lack of homing and cleavage in males developed above, we wanted to further optimize the system to ensure better drive performance. To do this we varied Cas9 expression and timing to test for improved homing efficiencies. The best performing *w^U6b-GDe^*/+ line was outcrossed to additional previously developed Cas9 strains (58), each with a unique promoter expressing Cas9: *4nitro-Cas9* (*4-nitrophenyl phosphatase*, AAEL007097), *trunk-Cas9* (trunk, AAEL007584), *nup50-Cas9* (nucleoporin 50kDa, AAEL005635), or *ubiq-Cas9,* which enabled us to test the effect of differential Cas9 expression on homing efficiency. Homing and cleavage efficiencies were evaluated in the G_1_ trans-heterozygotes *w^U6b-GDe^/w+*; *Cas9/+* with maternally or paternally inherited Cas9 (Fig. 2). The G_1_ trans-heterozygotes resulting from outcrosses of these maternal Cas9 lines to *wt* had a somatic mosaic *white* phenotype and cleavage rates of 19.4% ± 2.7% (*4nitro-Cas9)*, 83.9% ± 7.7% (*trunk-Cas9),* and 100% (*nup50-Cas9* and *ubiq-Cas9*) (Fig. 3D, Table S10). Moreover, zygotic Cas9 activity induced the somatic mosaic *white* phenotypes 49.4% ± 5.2% of the time in G_1_ trans-heterozygotes from paternally inherited *nup50-Cas9* crosses (Fig. 3D, Table S11). Interestingly, homing efficiencies varied by Cas9 strain; for example, trans-heterozygous *w^U6c-GDe^/w+*; *nup50-Cas9/+* females induced this somatic mosaicism in 100% ± 0% of their G_2_ progeny with an average 80.8% ± 7.9% inheritance rate in G_2_ progeny, while 49.2% ± 4.0% of G_2_ progeny from *w^U6c-GDe^/w+*; *nup50-Cas9/+* male crosses had this somatic mosaicism and 67.3% ± 5.2% of the G_2_ progeny inherited the drive (Fig. 2E–F; Tables S12–S13). The role of maternal effects in differential drive transmission was assessed by repeating these drive experiments with paternal Cas9 lines in G_2_, however we did not detect significant differences in drive inheritance between maternal and paternal Cas9 lines (*t*-test with equal variance, *P* > 0.05; Fig. 2E–F; Tables S14–S15).

### Multi-generation split-drive stability

Gene drives need to have a predictable behavior and stability across many generations, so these properties of the split-drive were determined over multiple consecutive generations by bidirectional outcrosses of *w^U6b-GDe^/w+; exu-Cas9/+* and *w^U6b-GDe^/w+; nup50-Cas9/+* females and males to *wt* lines, which should identify generational variations in cleavage or homing efficiencies indicating instability. In general, the cleavage and transmission rates were relatively stable over multiple generations (Fig. S6). However, super-Mendelian transmission of *w^U6b-GDe^* varied between individual mosquito crosses in both split-drive systems. For example, the average transmission rate for *w^U6b-GDe^/w+; exu-Cas9/+* females was 69.4% ± 8.5% (range 50–83%) and for *w^U6b-GDe^/w+; nup50-Cas9/+* females was 80.8% ±7.9% (range 59–94%); (Fig. S6; Tables S16–S17). Moreover, *w^U6b-GDe^/w+; exu-Cas9/+* males did not induce somatic *white* cleavage nor bias *w^U6b-GDe^* transmission (50.0% ± 0.5%; Fig. S6A, Table S18), while *w^U6b-GDe^/w+; nup50-Cas9/+* males did induce *white* cleavage in 49.2% ± 4.0% of progeny and biased transmission of the *w^U6b-GDe^* allele to 67.3% ± 5.2% of progeny over multiple generations (Fig. S6B; Table 19).

### Discovery of drive-resistant alleles

While tracking drive stability over successive generations, we consistently discovered drive-resistant alleles, which could limit the duration of spread of these drives in real-world settings. For example, *w^U6b-GDe^/w+; nup50-Cas9/+* females transmitted *white* mutant phenotypes to 0.31% ± 0.07% of progeny that did not inherit both split-drive transgenes, indicating that drive-resistant mutations were indeed generated at low frequency (Table S17). To determine whether these mutations occurred in the germline, we outcrossed these individuals to *white* recessive mutants, expecting to see white phenotypes if even one copy of *white* was disrupted in the germline. These experiments yielded 100% white-eye progeny, indicating that trans-heterozygous *w^U6b-GDe^/w+; nup50-Cas9/+* females deposited the Cas9 protein loaded with gRNA^w^ (Cas9/gRNA^w^) and this complex induced *white* germline mutations in progeny that did not inherit the *w^U6b-GDe^* transgene (Table S20). To confirm this on a molecular level, we performed genomic PCR/sequencing of the *white* locus from individual mutant mosquitoes and found variation in cleavage repair events (Fig. S7). For example, we identified loss-of-function resistance alleles (*w^R^*) that frequently contained indels at the *gRNA^w^* target site, making them unrecognizable to subsequent cleavage. This maternal carryover effect of the Cas9/gRNA^w^ complex was previously described in other drive systems and will likely be an issue when maternal promoters are utilized for drives (34, 43, 52, 59).

### Multi-release of split-drives results in high-frequency spread

Our next goal was to evaluate weekly releases of *Ae. aegypti* males homozygous for the split drive to determine whether this system could effectively drive an anti-pathogen gene (linked to the gRNA allele) into a population at a high frequency (≥80%) for an extended duration sufficient for disease control. To do this, we used our molecular work to inform the parameters (Table S22) of simulated split-drive releases of the best-performing system described above (*w^U6b-GDe^/w+; nup50-Cas9/+*). Modeling results suggested that 10 consecutive weekly releases of 10,000 homozygous split-drive males into a population with an equilibrium size of 10,000 adults would be sufficient to drive the split drive to a high frequency in the population (∼95% having at least one copy of the gRNA/anti-pathogen allele) (Fig. 4A). Mosquitoes with the gRNA/anti-pathogen alleles would then be slowly eliminated, falling to a frequency of ∼80% within four years of the final release. The inheritance bias induced when the Cas9 and gRNA co-occur would give the split-drive system an advantage over a comparable inundative release of mosquitoes with the anti-pathogen allele (Fig. 4B). A hypothetical construct with linked Cas9 and gRNA components and identical homing and fitness cost parameters demonstrated substantial spread after a single release of 10,000 adult males homozygous for the construct, but was outnumbered by in-frame resistant alleles three years following the release (Fig. 4C).

**Fig. 4.**
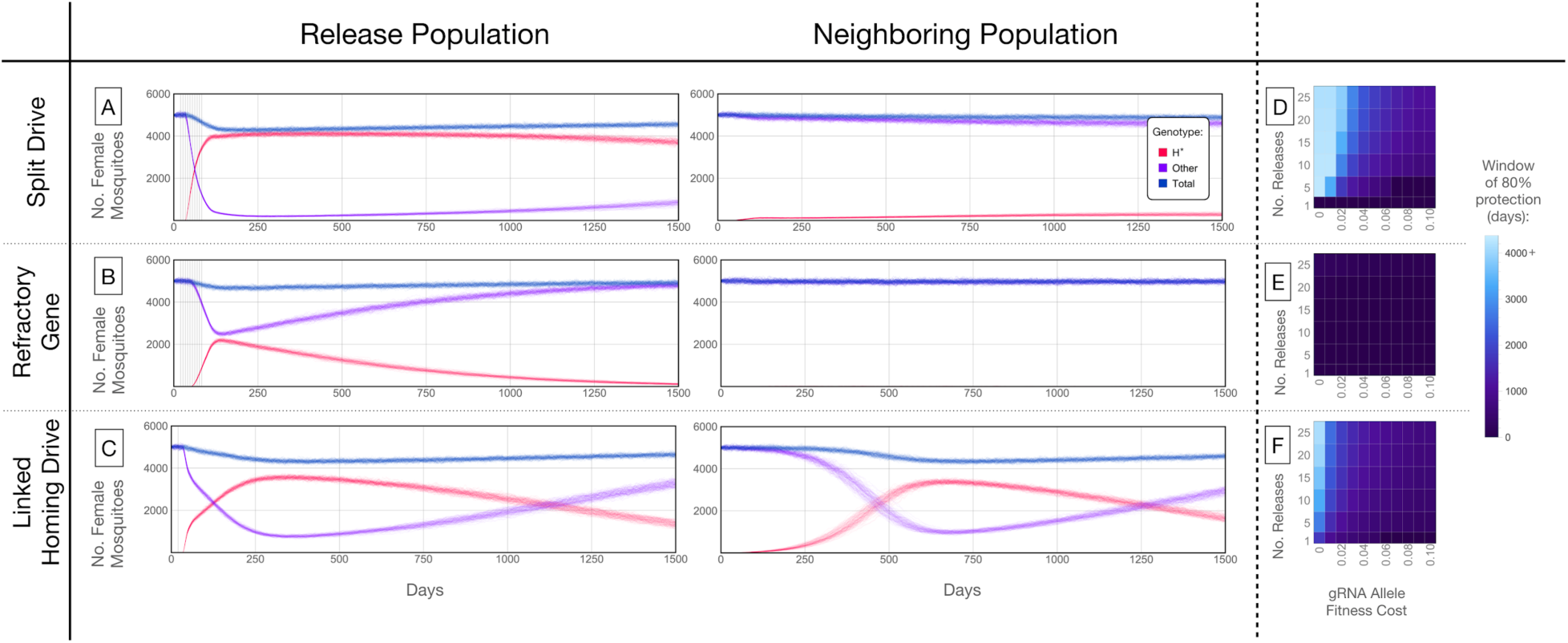
Mathematical model predictions for best performing split drive. Model predictions for releases of *Ae. aegypti* mosquitoes homozygous for (A) the split-drive system, (B) a disease-refractory gene, or (C) a homing drive system in which the components of the split-drive system are linked at the same locus (*C*). Parameters correspond to those for the best performing split-drive system (*w^U6b-GDe^/w*+*; nup50-Cas9/*+) (Table S22). Releases are carried out in a population with an equilibrium size of 10,000 adults and a 1% per mosquito per generation migration rate with a neighboring population of the same equilibrium size. Model predictions were computed using 100 realizations of the stochastic implementation of the MGDrivE simulation framework (79). Weekly releases of 10,000 homozygous split-drive males or the disease-refractory gene were simulated over a 10-week period, while a single release was simulated for the linked homing drive system. Total female population size (“total”, dark blue), adult females with at least one copy of the disease-refractory allele (“H*”, red), and disease-susceptible adult females without the disease-refractory allele (“other”, purple) were plotted for each group. Notably, the split-drive system is: i) largely confined to its release population, ii) reversible, and iii) present at a high frequency (>80% of adult females having at least one copy) for over four years. The split-drive system outperforms inundative adult male release of the disease-refractory gene in population disease refractoriness and outperforms the confinability of a linked homing drive system. In the right column, heatmaps are shown for (D) the split-drive system, (E) inundative disease-refractory gene, and (F) linked homing drive system and depict the window of protection in days that the proportion of mosquitoes in the release population with at least one copy of the disease-refractory allele exceeds 80%. The fitness cost (reduction in mean adult lifespan) per gRNA/refractory allele is varied along the x-axis, and the number of weekly releases along the y-axis. Notably, for the split-drive system, the window of protection exceeds four years following 10 or more weekly releases for gRNA/refractory allele fitness costs of 10% or less per allele.

### Split-drives are persistent and confineable

A key strength of a split-drive system is that it is expected to be reversible and largely confinable to a partially isolated population, making it of particular interest for field trials. As a comparison, the hypothetical linked homing drive discussed above, assuming a standard migrant exchange rate of 1% per mosquito per year, spreads to an over 50% allele frequency in the neighboring population following a single release (Fig. 4C, S9C). In contrast, the split-drive cargo (gRNA/anti-pathogen allele) reaches a peak frequency of ∼15% in the neighboring population four years after 10 releases (Fig. S9A) before being gradually eliminated by virtue of the fitness cost. In fact, this fitness cost causes the Cas9 allele to be eliminated in both populations, falling to a frequency of ∼14% in the release population within four years post release and barely reaches a frequency of 3% in the neighboring population, followed by a progressive decline of the gRNA/anti-pathogen allele in both populations. The reversal of the split-drive system can be accelerated by the release of wild-type males. After just 10 releases, the frequency of the split drive system in the population is expected to be sufficient for interrupting local disease transmission over a four year period. Fig. 4D depicts the duration for which the anti-pathogen allele is predicted to remain in over 80% of female mosquitoes when the number of releases and fitness cost of the gRNA/anti-pathogen allele are varied. This duration exceeds four years for the release scheme described above and for fitness costs less than 10% per allele.

## Discussion

*Ae. aegypti* is a globally distributed vector which spreads deadly pathogens to millions of people annually. Current control methods are inadequate for this vector and therefore new innovative technologies need to be developed and implemented. With the aim of providing new tools for *Ae. aegypti* control, here we engineer and test the first gene drives developed in this species. Importantly, to make these drives safe and confienable, we engineered these systems as split gene drives. These drives functioned at very high efficiency and may provide a tool to control this vector. To develop these drives, initially we functionally characterized several drive components that performed with high efficiency in both somatic cells and the germline. Additionally, we discovered a drive target sequence, within the *white* gene, that is highly conserved across many geographically distinct populations, providing a useful phenotypic readout of drive functionality, and demonstrating that highly conserved drive target sites can be found through extensive genomic sequencing of wild populations. Multiple bidirectional genetic crosses of transgenic lines with optimized drive components revealed that homing and cleavage efficiencies are largely dependent on the timing and expression of drive components and maternal deposition can result in embryonic activity and the generation of drive resistant alleles. Despite the creation of resistant alleles, however, mathematical modeling predicts that these split drives can spread locally and are also confinable and reversible, making them suitable for field trials and capable of local disease control once effectors are linked.

Notably, components characterized in this study including the polymerase III U6 promoters crossed with robustly Cas9 expression strains, can also be used to accelerate basic functional gene analysis in *Ae aegypti.* For example, tissue specific Cas9 strains could be crossed to U6-driven gRNA strains disrupting essential genes, to enable biallelic mutations in specific tissues (e.g. neurons) which otherwise would not be possible due to leathlity if disrupted in all cells. Additionally, if linked, these components could generate a potent drive that could self-disseminate for wider regional mosquito control. Our models predict that if these components were directly linked, assuming homing and cleavage rates similar to when unlinked, the drive would provide a window of >80% disease resistance among the female *Ae. aegypti* population for several years following a release (Fig. 4F) and spread from one population to another (Fig. 4C). These components could also be used in the future to develop self-limiting population suppression technologies, such as a precision-guided sterile insect technique (52), or to target conserved sex-determination genes such as doublesex (*dsx*) to generate a potent homing-based population suppression drive similar to the drive recently developed for *Anopheles gambiae* (*41*). Alternatively, these components could be used to generate a novel population suppression design by encoding a functional copy of *Nix* (56, 57), as the cargo, thereby converting females into fertile males while the drive spreads through and suppresses the population. Finally, these components support the development of other drives, including *trans*-acting toxin- and *cis*-acting antidote- (TA) based drives (52, 60), evolutionarily stable homing-based replacement drives that target haplosufficient essential genes and encode a recoded (i.e., cleavage resistant) rescue of the target gene (21, 52) or even TA underdominance-based drives (21), all of which could be instrumental for spreading anti-pathogen effectors (e.g., (26)) into wild disease-transmitting populations.

While this is a major step towards developing field-worthy gene-drive systems in *Ae. aegypti*, there are still limited effectors available to link with population modifying drives that will limit the immediate use of these technologies. The antiviral effectors developed in *Ae. aegypti* to date rely upon small RNA pathway-directed targeting of viral genomes (26, 61–64); however, these antiviral transgenes are not easily multiplexed and the diversity of arboviruses transmitted by this vector as well as the high mutation rate of these viruses (65) makes their practicality in the field tenuous. Additionally, long-term studies of viral resistance to these antiviral effectors are limited, so their long-term effectiveness is unknown. Furthermore, effector-associated fitness costs could impact the success of these technologies. While drive technologies should overcome some of these fitness costs, as we balance the optimization of drive efficacy with the minimization of drive risks, we will likely still need to develop effectors with reduced fitness costs. Current effectors have achieved efficacy against only a limited number of viral families and strains, usually those which are most highly lab adapted, and may have concerning fitness costs. Therefore, additional resources are needed to develop antiviral effectors that have reduced fitness costs, achieve broad antiviral activity against the diversity of viruses vectored by *Ae. aegypti,* and limit viral resistance despite high viral mutation rates.

It should be pointed out that gene-drive technologies are imperfect. Drive efficiencies are impacted by target-site availability, timing, the expression of drive components, host-specific factors such as the timing of meiosis and recombination, as well as the generation of resistant alleles that form as a result of non-homologous end-joining (NHEJ) repair of CRISPR-mediated DNA cleavage as opposed to homology directed repair (HDR). Notwithstanding, significant efforts are underway to optimize next-generation drives that incorporate molecular design architectures predicted to mitigate the generation and selection of resistant alleles and improve conversion efficiencies. These efforts include rational target-site design, regulation of the expression and timing of the nuclease, multiplexing drive targets, improved regulatory elements, and targeting ultra-conserved regions in essential genes that are recorded in the drive, enabling the selective elimination of non-functional NHEJ-repaired alleles (21, 52, 60, 66–68). While these future drive designs are expected to be more efficient than current systems, it should be noted that even imperfect drives with modest homing efficiencies are still predicted to be quite invasive (69), and will likely be useful for field implementation which could confer long-lasting entomological and epidemiological impacts.

Given the remarkable progress in the gene drive field and potential of gene drives to spread globally, understandably the discussion of the ethics, risks, governance, and the guidance of gene drives in the field have come to the forefront (44, 70–72). As gene-drive technologies advance and field trials become a necessary step toward understanding the efficacy and behavior of drives in wild populations, it is important that facilities planning and management, infrastructure upgrades, talent development for deployment and monitoring, and public engagement are advanced to support the sustainability of gene drive technologies. Safe, non-invasive, self-limiting split-drive technologies, such as the systems developed here, are a responsible choice for studying gene drives in the wild, as they safeguard against spreading to non-target populations and will allow the assessment of potential risks and unintended consequences. This may help establish the necessary infrastructure, while simultaneously controlling disease burdens, in preparation for a potential subsequent release of an efficient linked-drive system that can self-disseminate catalytically into broader landscapes. Taken together, our results provide compelling evidence for the feasibility of future effector-linked split-drive technologies that can contribute to the safe, sustained control and potentially the elimination of pathogens transmitted by this species.

## Materials and Methods

### Insect rearing

Mosquitoes used in all experiments were derived from the *Ae. aegypti* Liverpool strain (designated as wild-type [*wt*]). Of note, the *Ae. aegypti* reference genome sequences were generated using this strain (54, 73). Mosquitoes were raised in incubators at 28.0°C with 70–80% humidity and a 12-hour light/dark cycle. Larvae were fed ground fish food (TetraMin Tropical Flakes, Tetra Werke, Melle, Germany) and adults were fed 0.3 M aqueous sucrose. Adult females were blood fed three to five days after eclosion using anesthetized mice. Mosquitoes were examined, scored, and imaged using the Leica M165FC fluorescent stereo microscope equipped with the Leica DMC2900 camera. All animals were handled in accordance with the Guide for the Care and Use of Laboratory Animals as recommended by the National Institutes of Health and supervised by the local Institutional Animal Care and Use Committee (S17187).

### Profile of natural variation at *white* target site

To assess the natural variation within the *white* target site, we used data from individual whole genome sequencing of 133 *Ae. aegypti* specimens sampled in California (N = 123), Florida (N = 4), Mexico (N = 3), and South Africa (N = 3) (Fig. S1, Table S1) and sequenced to an approximate depth of 10x on an Illumina HiSeq 4000 producing 15.9 giga base pairs (Gbp). Then 15.5 Gbp of sequence reads were mapped with BWA-MEM v0.7.15 (74) to the latest version of the reference genome of *Ae. aegypti* (AaegL5 (54)). Polymorphisms were called with Freebayes v1.0.1 (75), applying all the default parameters except for ‘theta = 0.01’ and ‘max-complex-gap = 3’. We employed the most conservative approach by applying no filtering to the called polymorphisms to maintain the broadest possible set of potential variation. Additionally, we checked the polymorphism data for *Ae. aegypti* available on https://www.vectorbase.org (accessed 1/2/19) and found no published SNP’s in the region of the *white* target site. We therefore consider the target site very likely fixed in natural populations.

### Construct design and assembly

The Gibson enzymatic assembly method was used to build all constructs. To generate the gRNA constructs, the *piggyBac* plasmid (76) was digested with Fse1 and Asc1, and the linearized fragment was used as a backbone for construct assembly. Constructs for the U6 promoter screen contained the following fragments: the 5’-end flanking sequence of the U6 snRNA, which was selected as the promoter region; a 20-base sequence of the *gRNA^w^* (Fig. S2A); a 76-base fragment with a gRNA scaffold; and a 3xP3-eGFP-SV40 fragment. The first three fragments were generated using commercial gene synthesis (gBLOCK by IDT^®^ and GenScript). The 3xP3-eGFP fragment was amplified from a previously described plasmid harboring the 3xP3 promoter and coding sequence of eGFP (AddGene #111083 and 100705) using primers AE01, AE02, AE03, and AE04 (Table S21). A total of 12 plasmids were generated and are referred to as follows: U6a-T, U6b-T, U6c-T, U6d-T, U6e-T, U6f-T, U6g-T, U6h-T, U6i-T, U6j-T, U6k-T, and U6l-T (Fig. S2C).

We engineered four gene-drive element (GDe) constructs carrying different U6 promoters for site-specific integration at the *white* locus: *U6a-GDe*, *U6b-GDe, U6c-GDe*, and *U6d-GDe*. Each plasmid contained the following fragments (Fig. S2D): (1) left and right homology arms of ∼1 kb in length, which are complementary to the *Ae. aegypti white* locus immediately adjacent to the 5’ and 3’ ends of the *gRNA^w^* cut site, respectively, were gene synthesized by GenScript; (2) a U6 promoter with a 20-base *gRNA^w^* sequence and a 76-base gRNA scaffold, which was PCR amplified from U6a-T, U6b-T, U6c-T, and U6d-T constructs with the following primers: AE05 and AE06, AE07 and AE08, AE09 and AE10, AE11 and AE12, respectively; (3) the 3’UTR region of the corresponding U6 snRNA was gene synthesized by GenScript; (4) a 3xP3-tdTomato-SV40 fragment, which was amplified from the previously described plasmid (addgene #100705) using the following primers: AE13 and AE14, AE15 and AE14, AE16 and AE14, AE17 and AE14; (5) an Opie2-dsRed-SV40 fragment, which was amplified from the previously described plasmid (addgene #111083) using primers AE18 and AE19 (Table S21). All plasmids were grown in JM109 chemically competent cells (Zymo Research #T3005) and isolated using Zyppy^TM^ Plasmid Miniprep (Zymo Research #D4037) and Maxiprep (Zymo Research #D4028) kits. Each construct sequence was verified using Source Bioscience Sanger sequencing services. A list of primer sequences used in the above construct assembly can be found in Table S21. We have also made all plasmids and sequence maps available for download and/or order at Addgene (www.addgene.com) with identification numbers listed in Fig. S2.

### Embryo microinjection and mutation screening

Embryonic collection and CRISPR microinjections were performed following previously established procedures (77, 78). The concentration of plasmids used for the U6 promoter screen was 300 ng/µL. Injected G_0_ and G_1_ progeny were visualized at the larval, pupal, and adult life stages under a dissecting microscope (Olympus SZ51 and Leica M165FC). The heritable mutation rates were calculated as the number of G_1_ progeny with the loss-of-function mutation out of the number of all G_1_ progeny crossed with the white eye (*w–*) strain mosquitoes. To integrate each GDe construct at the *white* locus, a mixture containing 100 ng/µL of synthetic gRNA^w^, 100 ng/µL of each U6-GDe plasmid (U6a-GDe, U6b-GDe, U6c-GDe, and U6d-GDe), and 100 ng/µL of Cas9 protein was injected into 500 *wt* embryos for each plasmid. Synthetic gRNAs (Synthego) and recombinant *Streptococcus pyogenes* Cas9 protein (PNA Bio Inc.) were obtained commercially and diluted to 1,000 ng/µL in nuclease-free water and stored in aliquots at –80°C. A total of 233, 271, 191, 215 G_0_ adults were recovered for U6a-, U6b-, U6c-, and U6d-GDe injections, respectively. Successful integration into the *white* locus was determined by visually identifying the eye-specific 3xP3-tdTomato fluorescence in G_1_ heterozygous mosquito larvae with black eyes (*w^U6-GDe^/w+*) and in G_2_ homozygous mosquito larvae with white eyes (*w^U6-GDe^/w^U6-GDe^*). In addition, site-specific integration of U6-GDe constructs was confirmed by amplifying and Sanger sequencing both the left and right integration points (Fig. S5) from a genomic DNA prep of each *w^U6-GDe^* line with the following primers: AE20, AE21, AE22, and AE23 (Table S21).

### Assessment of split-gene-drive efficacy

Both gene-drive (*GD)* and *Cas9* elements are required for a functional split-gene drive. To generate G_1_ trans-heterozygous *w^U6-GDe^/w+; Cas9/+* male and female mosquitoes with an active split-gene drive, we reciprocally crossed *w^U6-GDe^/w+* and *Cas9/Cas9* strains. While selecting trans-heterozygous mosquitoes, we scored the loss-of-function mutation of *white* in all G_1_ progeny. Then, G_1_ trans-heterozygous mosquitoes were mated to *wt* for 4 days, and the *white* mutation and *w^U6-GDe^* transmission frequencies were calculated in G_2_ progeny. Females were blood fed on anesthetized mice on the fifth day. Three days later, more than twenty females were allowed to lay eggs individually into a 50-mL vial filled with water and lined with wet filter paper. The larval progeny from each female were counted and scored for *w^U6-GDe^* and *Cas9* by *3xP3-tdTomato* and *Opie2-dsRed* expression, respectively. Females that failed to take blood and produce progeny were excluded from the analysis. According to Mendelian inheritance, the expected G_2_ transmission rate of *w^U6-GDe^* is 50% under this crossing scheme; the higher rate indicates super-Mendelian inheritance (Fig. 2A). *White* mutagenesis and *w^U6-GDe^* transmission rates by trans-heterozygous *w^U6-GDe^/w+; Cas9/+* mosquitoes each carrying *GDe* with four different U6 promoters were compared to those of the corresponding *w^U6-GDe^/w+* line without *Cas9* in the genetic background. The homing and cleavage efficiencies of the two split-drive systems, *w^U6b-GDe^/w+; exu-Cas9/+* and *w^U6b-GDe^/w+; nup50-Cas9/+*, were observed for the following four generations by scoring the markers in the progeny from reciprocal crosses of trans-heterozygous and *wt* mosquitoes.

### Molecular characterization of *w^R^* alleles

To test whether *w^R^*-resistant alleles were present in germ cells, mosquitoes harboring *w^R^* alleles were genetically crossed into the *w^19Δ^/w^19Δ^* recessive mutant background, and their progeny was scored for *white* loss-of-function mutations. Genomic DNA was extracted from an individual mosquito of each progeny group using the DNeasy Blood & Tissue Kit (QIAGEN) following the manufacturer’s protocol, and a *white* region carrying the *gRNA^w^* target site was PCR amplified with primers AE24 and AE25. Because the tested mosquitoes were heterozygous *w^R^/w^19Δ^* for *white* alleles, a PCR amplicon from each mosquito was first cloned into an open plasmid using Gibson enzymatic assembly, and seven clones were PCR amplified and Sanger sequenced to identify novel *w^R^* alleles. *De novo-*mutated *w^R^* alleles were identified by comparing them with both *w+* and *w^19Δ^* alleles using Sequencher^TM^ and 5.0 SnapGene® 4.2. The same method was used to confirm the presence of *w+* non-cleaved alleles in a few mosquitoes with black eyes, which were later inferred to be *w+/w^19Δ^* and *w+/w^U6b-GDe^*.

### Mathematical modeling

To model the expected performance of a split-drive functioning as a confinable and reversible gene-drive system and as a test system prior to the release of a linked homing drive for *Ae. aegypti*, we simulated release schemes for split-drive, linked homing drive, and inundative releases of males carrying a refractory allele using the MGDrivE simulation framework (79) (https://marshalllab.github.io/MGDrivE/). This framework models the egg, larval, pupal, and adult mosquito life stages (both male and female adults are modeled) implementing a daily time step, overlapping generations, and a mating structure in which adult males mate throughout their lifetime while adult females mate once upon emergence, retaining the genetic material of the adult male with whom they mate for the duration of their adult lifespan. Density-independent mortality rates for the juvenile life stages are assumed to be identical and are chosen for consistency with the population growth rate in the absence of density-dependent mortality. Additional density-dependent mortality occurs at the larval stage, the form of which is taken from previous studies (*80*). The inheritance patterns for the split-drive, linked homing drive, and refractory gene systems are modeled within the inheritance module of the MGDrivE framework (79) along with their impacts on female fecundity and adult mortality rate. We parameterized our split-drive model according to the best-performing system in this study (*w^U6b-GDe^/w+; nup50-Cas9/+*): i) a cleavage frequency of 100% in females and 49.2% in males, ii) HDR frequency given a cleavage of 61.6% in females and 70.3% in males, and iii) each Cas9 allele was associated with an ∼11.2% reduction in female fecundity (Fig. S8). Resistant allele generation rates were assumed from previously analyzed CRISPR-based homing constructs (36) (∼15% of non-HDR events lead to in-frame/cost-free resistant alleles, with the remaining leading to out-of-frame/costly resistant alleles). By default, we assumed a 5% reduction in mean lifespan associated with each copy of the gRNA/disease-refractory allele. We implemented the stochastic version of the MGDrivE framework to capture the randomness associated with rare events such as resistant allele generation. We simulated two partially isolated populations each consisting of 10,000 adults at equilibrium, exchanging migrants at a rate of 1% per mosquito per generation, with weekly releases of 10,000 adult males over a defined period. *Ae. aegypti* life history and intervention parameter values are listed in Table S22.

### Statistical analysis

Statistical analysis was performed in JMP 8.0.2 by SAS Institute Inc. Twenty replicates were used to generate statistical means for comparisons. *P* values were calculated for a two-sample Student’s t-test with equal variance. Point plots were built in Prism 7 by GraphPad.

## Acknowledgements

This work was supported by funding from a Defense Advanced Research Project Agency (DARPA) Safe Genes Program Grant (HR0011-17-2-0047) awarded to O.S.A. and subcontracted to G.C.L. and J.M.M. We acknowledge funding support from the UC Davis Bridge Funding Program, UC Davis School of Veterinary Medicine Vector-Borne Disease Pilot Grant Program, and the Pacific Southwest Regional Center of Excellence for Vector-Borne Diseases funded by the U.S. Centers for Disease Control and Prevention (Cooperative Agreement 1U01CK000516). We thank Judy Ishikawa for help rearing and maintaining all mosquito strains produced in this study. We also thank Allison Weakley, Kendra Person, and Hans Gripkey for field mosquito collection data processing including DNA extraction, DNA quantification, and library preparations. We thank personnel from Consolidated Mosquito Abatement District (Ms. Jodi Holeman and Ms. Katherine Ramirez), Coachella Valley, Delta, Greater LA County (Dr. Susan Kluh), and San Mateo County Vector Control Districts and Fresno, Kern, Madera County, Northwest (Dr. Major Dhillon, District Manager), Orange County (Mr. Michael Hearst, District Manager), and San Gabriel Valley Mosquito and Vector Control Districts, Kings Mosquito Abatement District, Community Health Division of the Department of Environmental Health (Ms. Rebecca Lafreniere, Chief and Ms. Elizabeth Pozzebon, Director), Imperial County Public Health Department, San Bernardino County Mosquito & Vector Control Program, San Diego County Dept. of Environmental Health, Vector Control, Dr. Chelsea Smartt (Florida Medical Entomology Laboratory), and Dr. Christopher Barker (UC Davis) for providing specimens used in this study. We also thank Dr. Anthony J. Cornel (UC Davis) and Dr. Leo Braack (University of Pretoria, South Africa) for the collection of South African samples and Dr. Danny Governer and SANParks for permitting collection from Shingwedzi in the Kruger National Park. We also thank Dr. Lutz Froenicke and his team at the UC Davis DNA Technologies Core for genome sequencing.

## Author Contributions

O.S.A., M.L., and N.P.K. conceptualized the study. T.Y. assembled all constructs for this project. M.L., M.B., and S.G. performed molecular and genetic experiments. R.R. contributed to the interpretation of the results and project management. Y.L. carried out whole genome sequencing of *Ae. aegypti* field isolates. H.S. performed resistant allele analysis. G.C.L. conducted field sample acquisition and contributed to genome sequencing and resistant allele analysis. H.M.S.C., J.B., and J.M.M. conducted the mathematical modeling. All authors contributed to the writing, analyzed the data, and approved the final manuscript.

## Supplementary Figures

**Fig. S1.**
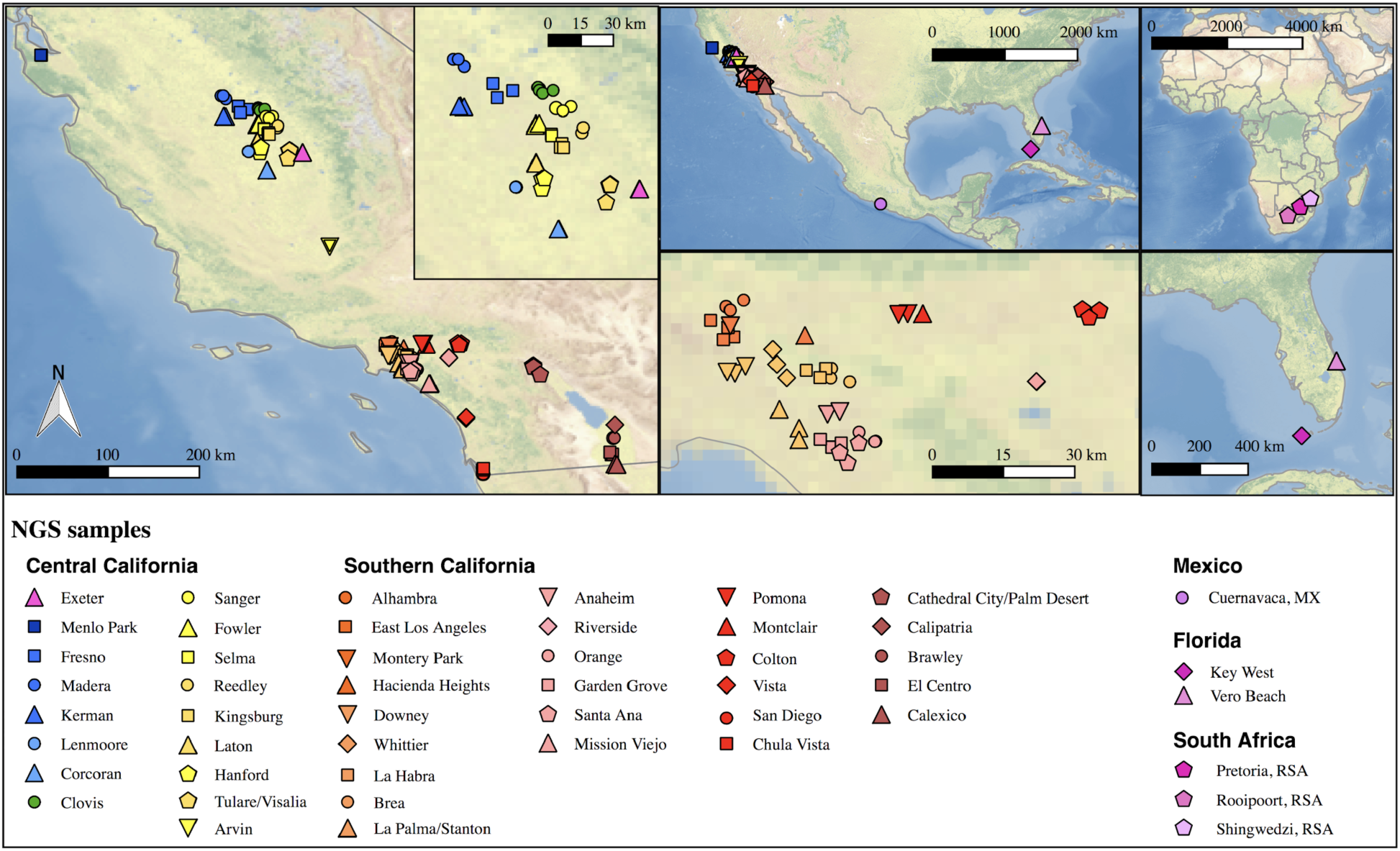
Sample locations of mosquitoes utilized for whole genome sequencing. Left map depicts the location of all samples from California (CA) with the inset enlarging the Fresno/Clovis area. Top center shows a North American map including CA, Florida, and Cuernavaca, Mexico. Bottom central shows southern California near Greater Los Angeles area. Top right shows an African map with three locations in South African sample origin. Bottom right shows a Florida map with sampling locations. Each symbol represents a single individual mosquito. These plots were generated using CleanTOPO2 basemap (81)

**Fig. S2.**
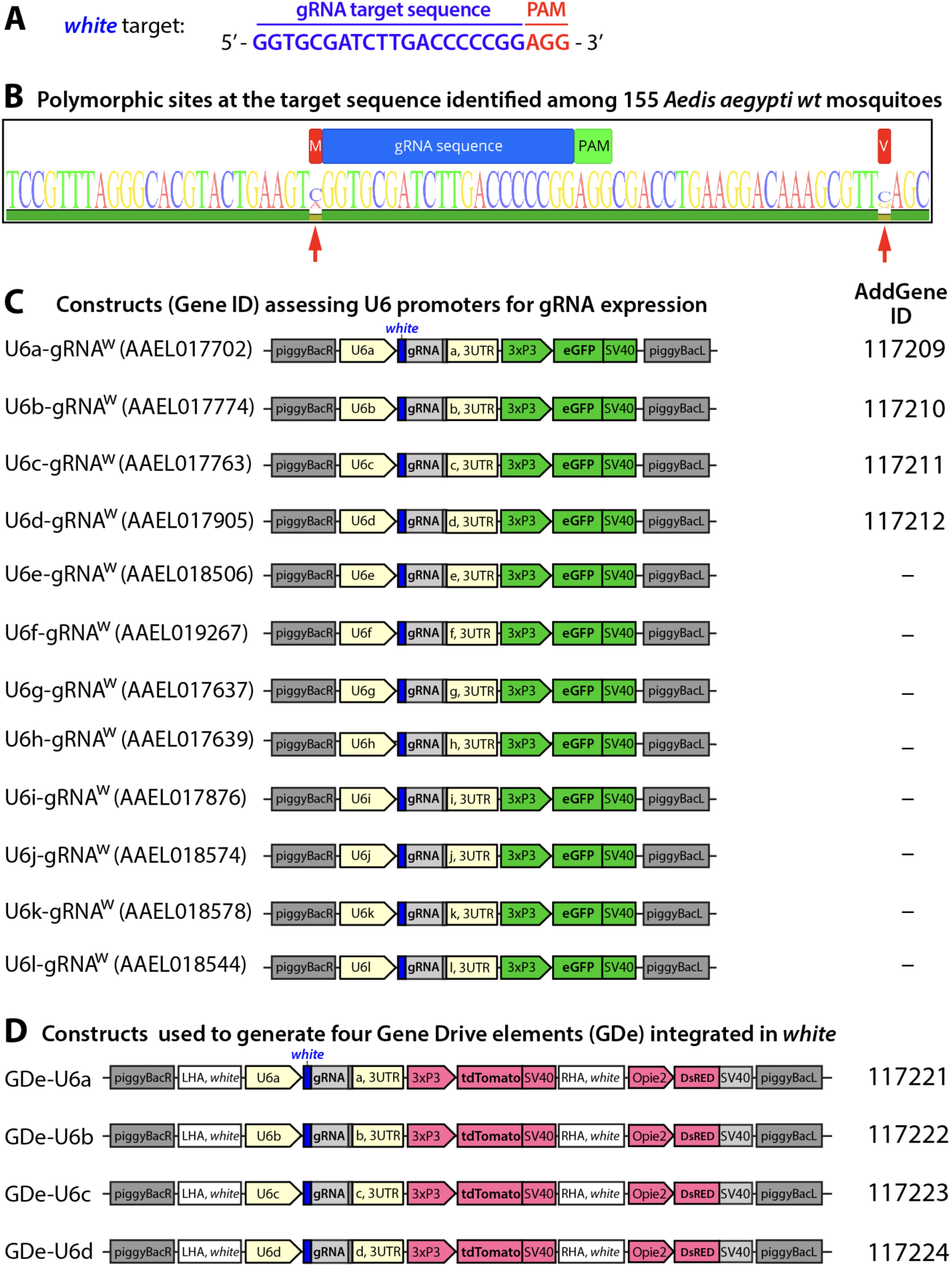
Guide RNA (gRNA) sequence and gene constructs developed in the study. Nucleotide sequence for the *gRNA^w^* genomic target sequence in exon 3 of *white* gene with the PAM sequence indicated in red (A). Conservation of the *RNA^w^* genomic target sequence identified among 133 *Ae. aegypti* mosquitoes sampled from the field (*B*). Within the target sequence (AaegL5. 1:107,955,440-107,955,462, Vectorbase.org) (55)(54) no polymorphism sites were detected. The nearest polymorphic sites detected (red arrows) were −1 bp upstream (107,955,439) and +22 bp downstream (107,955,484). Schematic maps of twelve constructs utilized to functionally test U6 promoter activity in *Ae. aegypti* (*C*). Schematic maps of the four split gene drive elements (*GDe*) developed in this study. Plasmids generated in the study were deposited at Addgene.org and the corresponding ID numbers are listed.

**Fig. S3.**
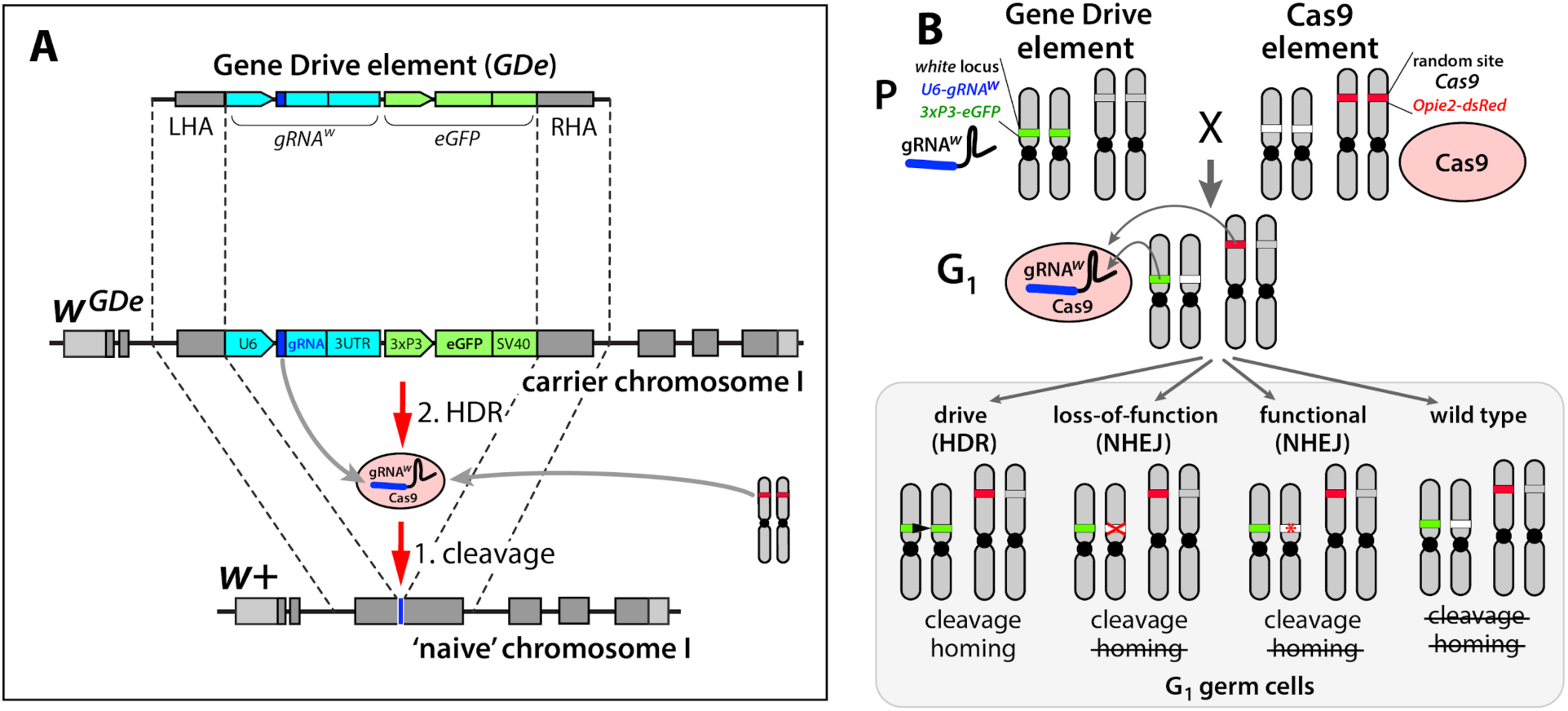
Design of the split-drive, a confined and high-threshold population replacement system. To confine a gene drive’s propensity to spread, we separated the complete CRISPR/Cas9 gene drive system into two components: a Gene Drive element (GDe) comprising a gRNA and a fluorescent marker both flanked by Left and Right Homology Arms (LHA and RHA); and Cas9 endonuclease (Cas9). GDe integrated in a target gene, here it was *white* locus, can spread from the carrier chromosome into a naive chromosome in the presence of *Cas9* transgene randomly integrated at a separate locus (*A*). First, Cas9/gRNA complex cuts the target site (cleavage); when GDe can be copied from the carrier chromosome into a naive chromosome via Homology Directed Repair (HDR). Each element is inactive on its own and can be maintained as a homozygous parental line (P). The cross between the homozygous lines results in 100% trans-heterozygous G_1_ progeny that carry both elements (*B*). gRNA expressed by GDe directs cleavage at *white* locus by Cas9, which can be repaired in three different ways: via HDR using GDe as a repair template and result in homing of GDe, and via Non-Homology End Joining (NHEJ) and lead to a loss-of-function mutant allele or a in-frame functional mutant allele. NHEJ-induced mutated alleles carry insertions/deletions (indels) at the cut site and become resistant to the same Cas9/gRNA system.

**Fig. S4.**
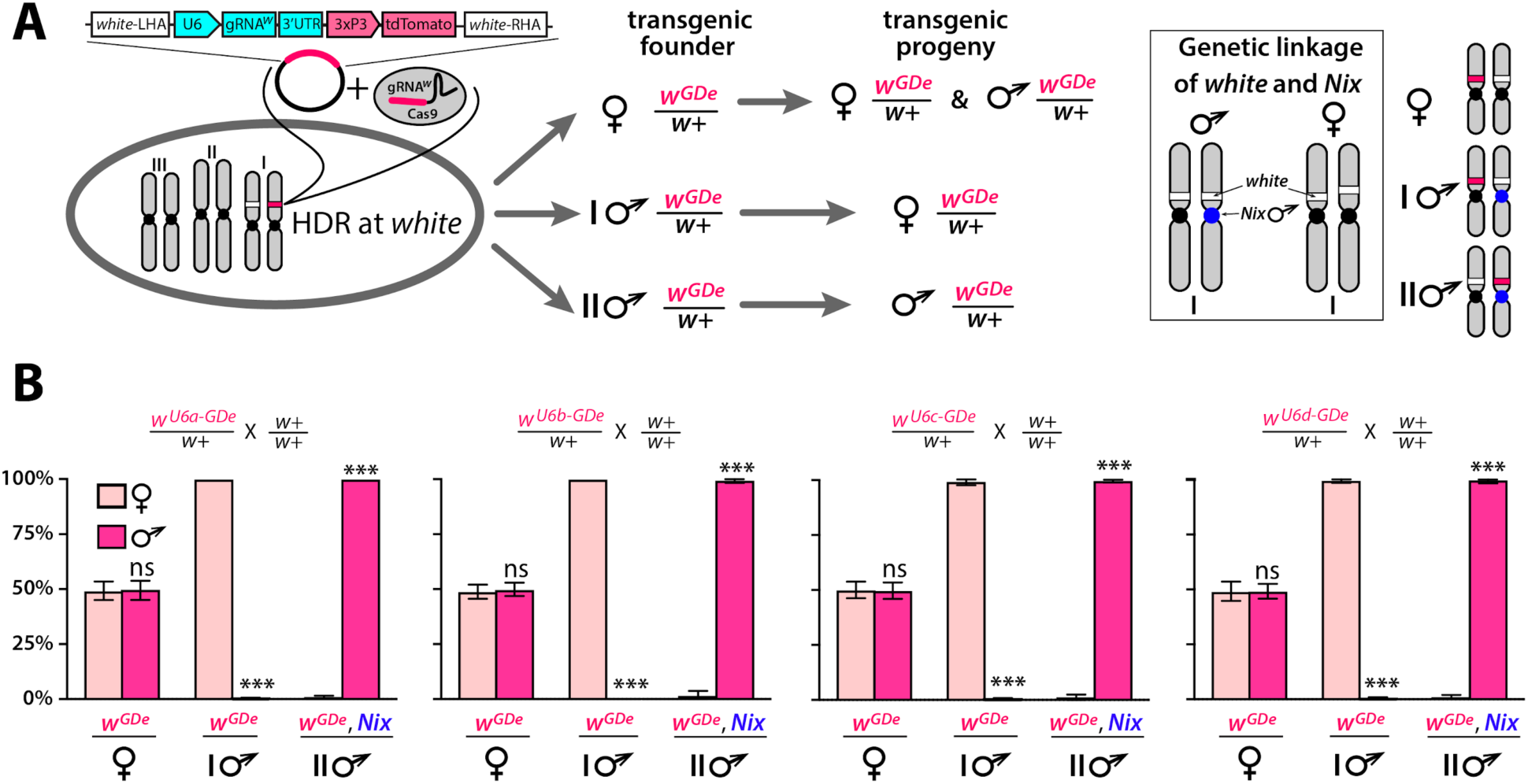
Sex biased inheritance of the GDe is due to linkage of *white* and *Nix*, a male determining gene in *Ae. aegypti*. Schematic of site-specific integration and sex-linked inheritance of Gene Drive element (GDe) (*A*). To direct site-specific integration of GDe inside *white* locus (white stripe on chromosome I) via homology directed repair (HDR), wild type (*wt*) *Ae. aegypti* embryos were injected with a Cas9/gRNA^w^ complex and a plasmid carrying GDe flanked by homology regions complementary to left and right genomic regions at the *white* cut site. GDe contains two genes: *U6-gRNA^w^* to direct a site-specific cleavage and the *3xP3-tdTomato* transgenesis marker (red stripe). Female mosquitoes carrying *w^GDe^/w+* passed *w^GDe^* randomly to both genders, while transgenic males transmitted *w^GDe^* either to females (type I) or males (type II) nearly exclusively. Close genetic linkage of *white* and *Nix* genes causes tight sex-linked inheritance of *w^GDe^* via male founders (*B*). *Nix* is located near the centromere of chromosome I (54, 55) and consequently, type I male (**♂**) founders have *w^GDe^* inserted on the chromosome copy without *Nix* and therefore pass *w^GDe^* exclusively to female progeny. On the other hand, type II male founders have *w^GDe^* integrated on the chromosome with *Nix*, and thus they transfer *w^GDe^* and *Nix* together exclusively to male progeny (box). Bars show average ± SD estimated for 20 data points. Statistical significance between gender frequencies was estimated by an equal variance *t* test. (*P* ≥ 0.05^ns^ and *P* < 0.001***).

**Fig. S5.**
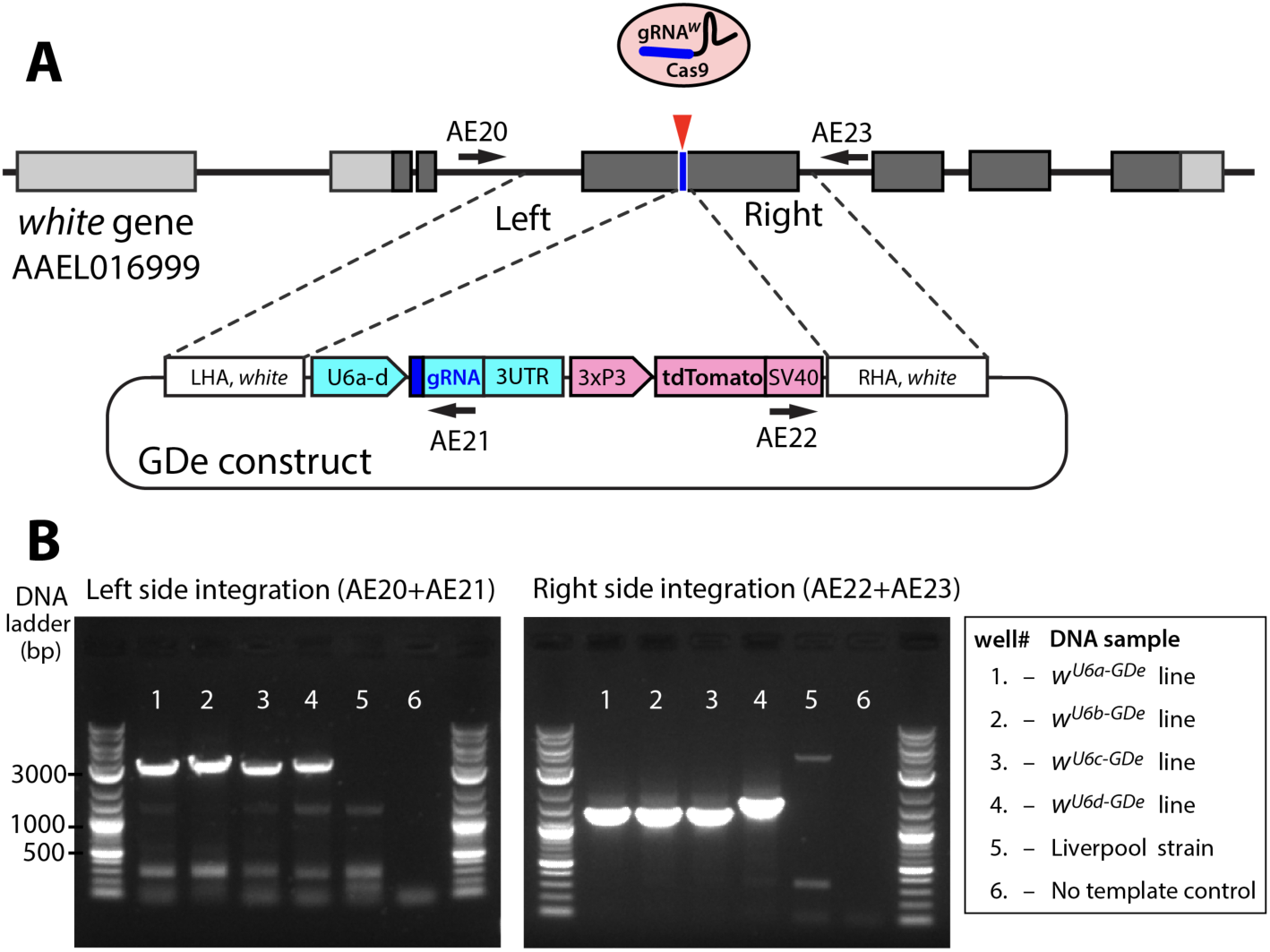
Site-specific integration of Gene Drive element (GDe). Schematic showing positions of Left and Right Homology Arms (LHA and RHA) of *GDe* relative to genomic *white* sequence. The cut site is presented over the genetic structure of *white,* with darker gray boxes representing exonic coding sequences. GDe constructs were integrated at the cut site via homology directed repair (HDR) relying on the construct’s LHA and RHA to *white.* Arrows depict primers (AE20, AE21, AE22 and AE23) used for PCR and their location relative to genomic and constructs sequences. Two gel images showing specific PCR amplicons for each side of integration (*B*). The same DNA samples, listed in the box to the left, were used to PCR of both fragments. To confirm PCR specificity, amplicons were Sanger sequenced from both ends.

**Fig. S6.**
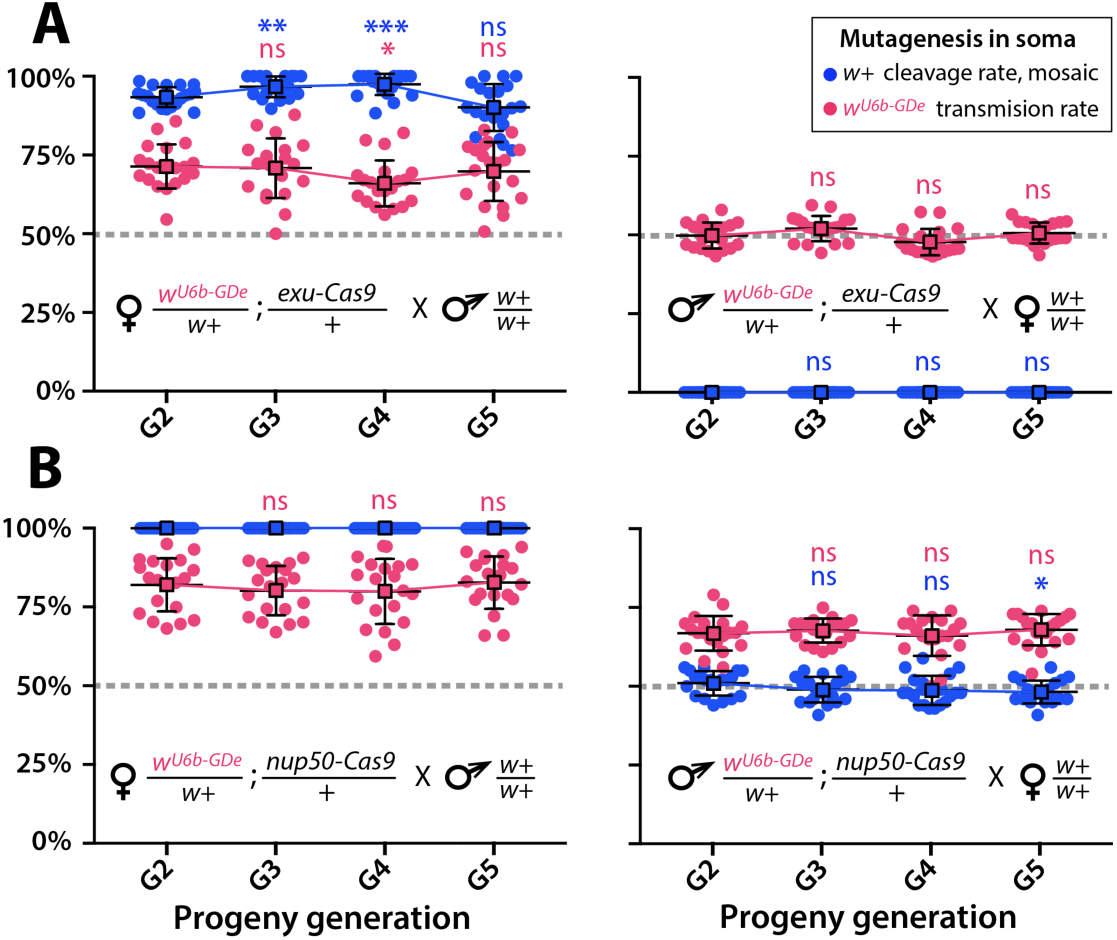
Split-drive over multiple generations. Frequencies of *white* cleavage and *w^U6b-GDe^* transmission were plotted over multiple generations. The *w^U6b-GDe^/w+; exu-Cas9/+* and *w^U6b-GDe^/w+; nup50-Cas9/+* trans-heterozygous females, or males, were outcrossed to *wt* each generation and both transmission and cleavage rates were scored. Average transmission rates at consecutive generations were compared to the corresponding rates at G_2_. Point plots show the average ± SD over 20 data points. Statistical significance was estimated using a *t* test with equal variance. (*P* ≥ 0.05^ns^, *P* < 0.05*, and *P* < 0.001***).

**Fig. S7.**
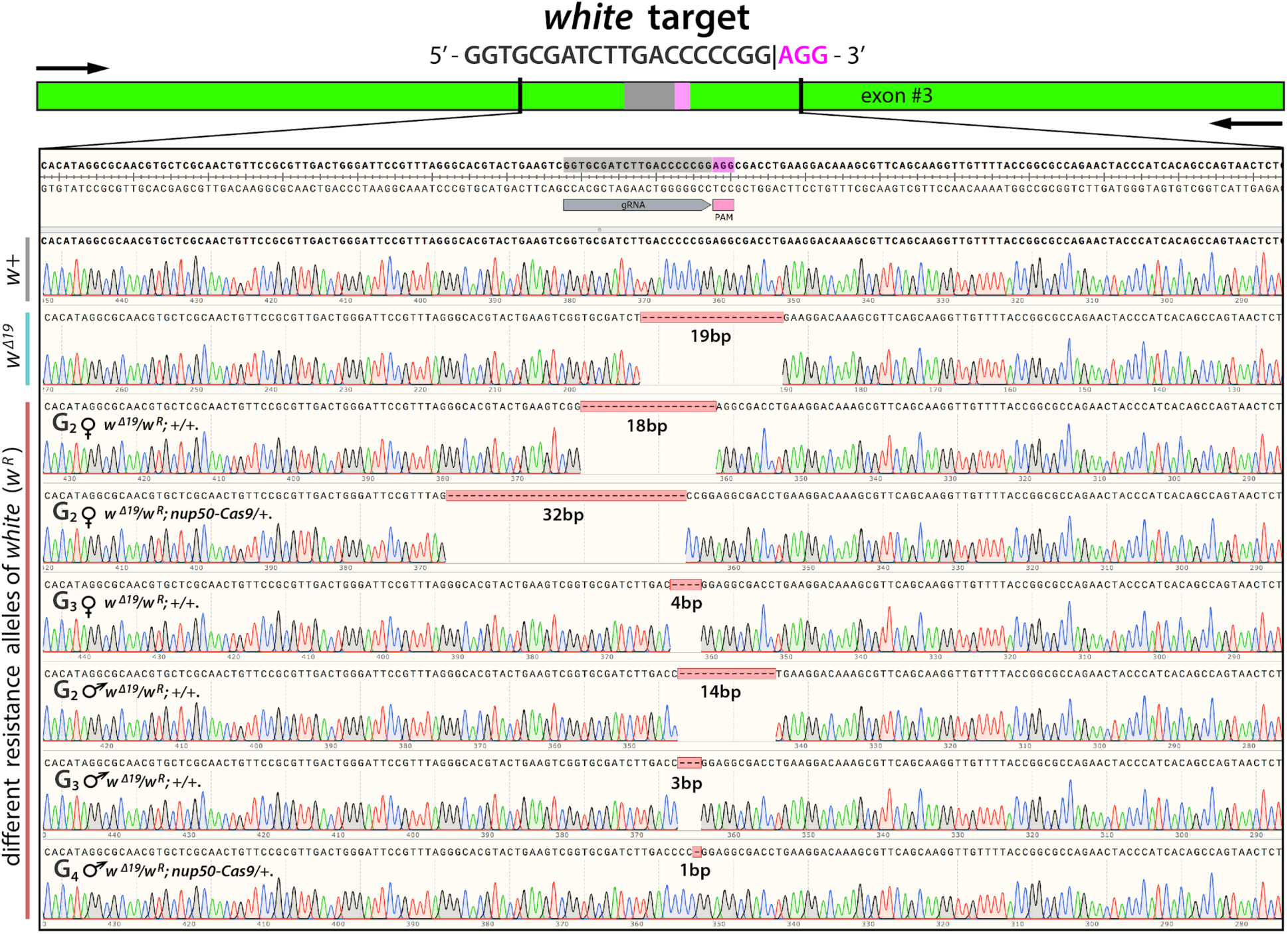
Sequences of *de novo* resistance alleles at *white* locus (*w^R^*). Mosquitoes carrying resistance alleles (*w^R^)* were identified among progeny of genetic crosses between *w^U6b-gRNA^/w+; nup50-Cas9/+* trans-heterozygous mosquitoes with *wt* (Table S20), and were further crossed to the *Ae. aegypti* loss-of-function *w^Δ19^/w^Δ19^* line, to place novel knockout alleles (*w^R^*) into the recessive genetic background. To sample *w^R^* alleles from *w^Δ19^/w^R^* heterozygous mosquitoes, PCR amplicons amplified from an individual mosquitoes were cloned into a plasmid, and a seven clones were Sanger sequenced in both directions and aligned against the *w+* and *w^Δ19^* alleles in SnapGene® 4.2. Genders and genotypes of analyzed mosquitoes as well as the generation when they were recovered are provided on the left side. Numbers of bases deleted is indicated below the corresponding deletion. Notably, while the deletions of 3 and 18 bases re-established the reading frame for *white* gene, they still displayed the loss-of-function phenotype.

**Fig. S8.**
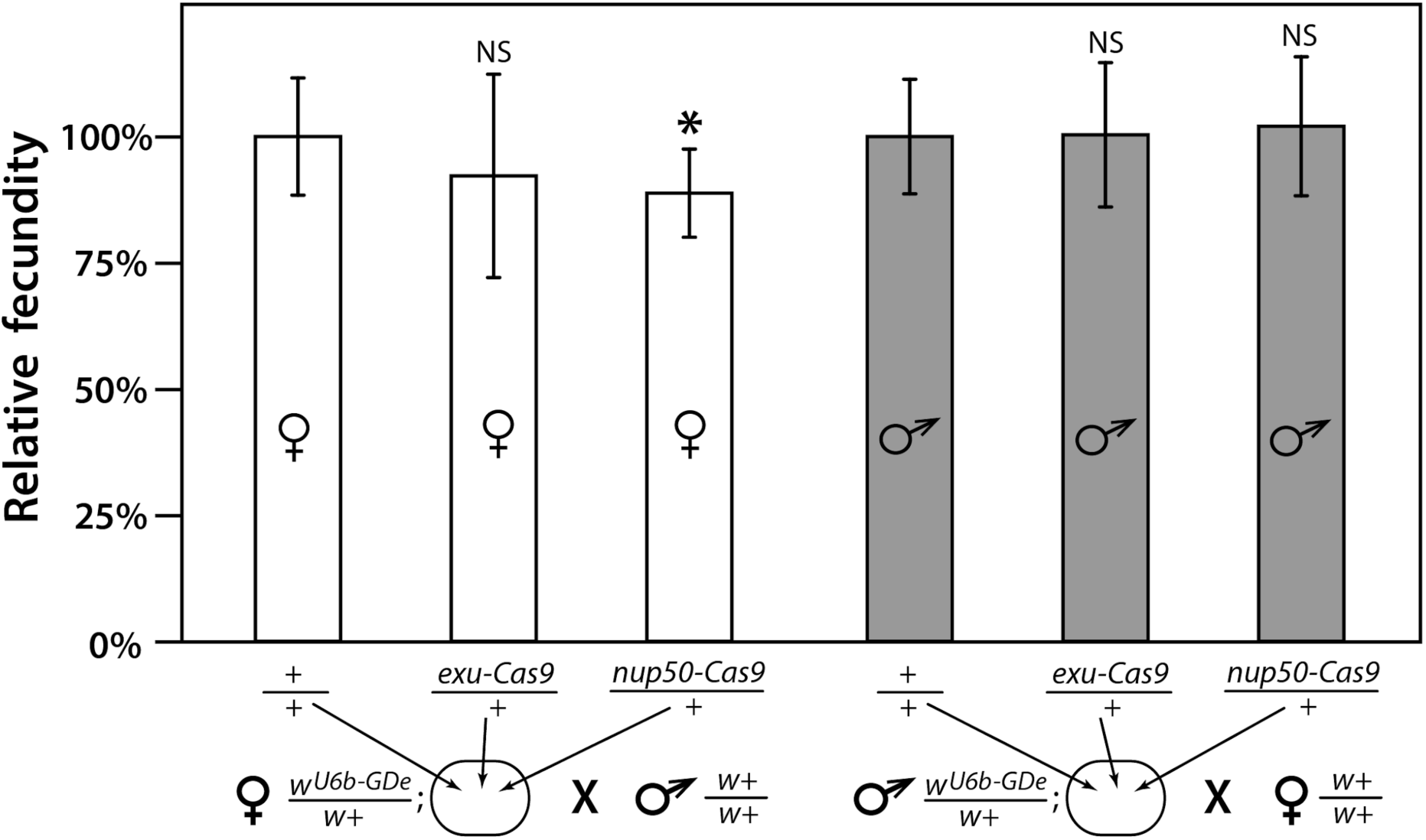
Fecundity of trans-heterozygous females only slightly reduced. Progeny numbers from 20 individual crosses for each parent combination crossed to *wt* (*w+/w+*) mosquitoes of the opposite sex were used to estimate fecundity. To transform progeny numbers into fecundity percentages, each progeny number was normalized to the average progeny number in the corresponding control group, heterozygous *w^U6b-GDe^/w+* females or males crossed to *w+/w+* mosquitoes. *nup50-Cas9* caused a significant reduction in fecundity, from 100.0% ± 11.6% to 88.8% ± 8.7%. Average fecundity percentages were compared to that in the corresponding control group. Bars show averages ± SD estimated from 20 data points. Statistical significance was estimated by a *t* test with equal variance. (*P* < 0.02*).

**Fig. S9.**
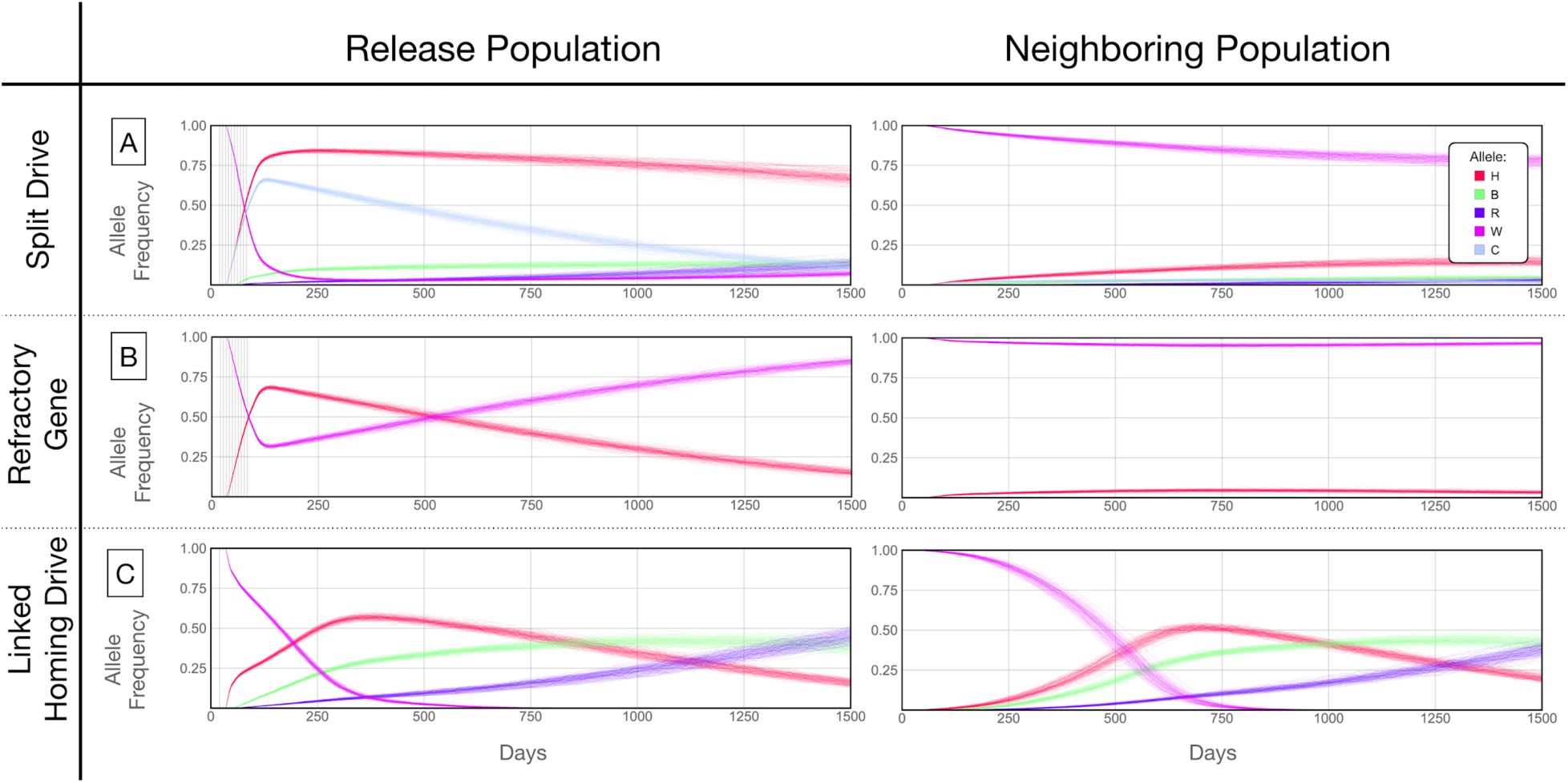
Allele frequencies for best performing split-drive construct. Model predictions for releases of *Ae. aegypti* mosquitoes homozygous for the split-drive system (*A*), a disease-refractory gene (*B*), or a homing drive system in which the components of the split-drive system are linked at the same locus (*C*). Parameters correspond to those for the best performing split-drive system (*w^U6b-GDe^/w+; nup50-Cas9/+*) (Table S22). Releases are carried out in a population with an equilibrium size of 10,000 adults that exchanges migrants with a neighboring population of the same equilibrium size at a rate of 1% per mosquito per generation. Model predictions were computed using 100 realizations of the stochastic implementation of the MGDrivE simulation framework (79). Weekly releases of 10,000 males homozygous for the split-drive system or disease-refractory gene were carried out over a 10 week period, while a single release was carried out for the linked homing drive system. Results are plotted for female allele frequencies. Red denotes the gRNA/disease-refractory allele (H), pink denotes the wild-type allele at this locus (W), and dark blue and green represent in-frame/cost-free and out-of-frame/costly resistant alleles (R and B). Light blue represents the allele frequency of Cas9 at the second locus for the split drive system (C). Notably, the H allele reaches a frequency of ∼15% in the neighboring population for the split-drive releases, before being eliminated by virtue of a fitness cost. At the release site, the Cas9 allele is reduced to a population frequency of <15% within four years of the final release, leading to a progressive decline of the gRNA/refractory allele in both populations. Reversibility can be accelerated through dilution of transgenes by releases of wild-type males.

## Supplemental Tables

**Table S22.** Parameter values used in *Aedes aegypti* population model.

| System | Parameter | Value | Reference |
| --- | --- | --- | --- |
| General | Egg production per female (day<sup>-1</sup>) | 20 | (82) |
|  | Duration of egg stage (days) | 5 | (83) |
|  | Duration of larval stage (days) | 6 | (83) |
|  | Duration of pupal stage (days) | 4 | (83) |
|  | Daily population growth rate (day<sup>-1</sup>) | 1.175 | (84) |
|  | Daily mortality risk of adult stage (day<sup>-1</sup>) | 0.090 | (85)(86) |
|  | Adult female population size | 10,000 | (87) |
|  | Migration rate (mosquito<sup>-1</sup> day<sup>-1</sup>) | 0.01 | (88) |
| Split or linked homing drive | Cleavage rate (female) | 1 | This paper |
|  | Cleavage rate (male) | 0.492 | This paper |
|  | Frequency of accurate homology-directed repair (HDR) given cleavage (female) | 0.616 | This paper |
|  | Frequency of accurate HDR given cleavage (male) | 0.703 | This paper |
|  | Proportion of non-HDR events that lead to in-frame/cost-free resistant alleles | 0.15 | (36) |
|  | Reduction in female fecundity per Cas9 allele | 0.112 | This paper |
| Homing drive or inundative release | Reduction in adult lifespan per gRNA/refractory allele | 0.05 |  |

